# Enhancer regulatory networks globally connect non-coding breast cancer loci to cancer genes

**DOI:** 10.1101/2023.11.20.567880

**Authors:** Yihan Wang, Daniel Armendariz, Lei Wang, Huan Zhao, Shiqi Xie, Gary C. Hon

## Abstract

Genetic studies have associated thousands of enhancers with breast cancer. However, the vast majority have not been functionally characterized. Thus, it remains unclear how variant-associated enhancers contribute to cancer. Here, we perform single-cell CRISPRi screens of 3,512 regulatory elements associated with breast cancer to measure the impact of these regions on transcriptional phenotypes. Analysis of >500,000 single-cell transcriptomes in two breast cancer cell lines shows that perturbation of variant-associated enhancers disrupts breast cancer gene programs. We observe variant-associated enhancers that directly or indirectly regulate the expression of cancer genes. We also find one-to-multiple and multiple-to-one network motifs where enhancers indirectly regulate cancer genes. Notably, multiple variant-associated enhancers indirectly regulate TP53. Comparative studies illustrate sub-type specific functions between enhancers in ER+ and ER- cells. Finally, we developed the pySpade package to facilitate analysis of single-cell enhancer screens. Overall, we demonstrate that enhancers form regulatory networks that link cancer genes in the genome, providing a more comprehensive understanding of the contribution of enhancers to breast cancer development.

## INTRODUCTION

Cancer sequencing studies have established the role of genes in cancer. While millions of non-coding genetic variants have been identified in cancer patients, only a handful have been functionally characterized with mechanistic links to cancer development ^1–3^. For example, non-coding somatic variants are enriched at mutation hotspots including the telomerase reverse transcriptase (TERT) promoter ^2,3^. Systematic functional analysis of variant-associated non-coding regions could improve our understanding of the genetic and molecular mechanisms of cancer.

Two approaches have played pivotal roles in understanding the genetic basis of cancer. Genome-wide association studies (GWAS) have identified thousands of single nucleotide variants (SNVs) associated with breast cancer by genotyping large patient cohorts ^4–9^. Since these studies sample DNA from normal rather than cancerous cells, GWAS variants represent genetic changes that contribute to cancer risk, rather than those that directly drive cancer development. To address the limitation of GWAS, whole genome sequencing (WGS) studies have pinpointed the somatic mutations that drive cancer by comparing the genomes of tumor cells and non-tumor cells from the same patient ^10–12^. The majority of genetic variants identified in GWAS and WGS are non-coding. Many of these variants contain epigenetic signatures of enhancers ^13–16^. Since enhancers bind transcription factors to orchestrate gene expression programs in cancer cells ^17–20^, one model is that non-coding variants modify the activity of transcriptional enhancers, thereby contributing to cancer development by altering the expression of cancer genes. Several published examples support this model. Michailidou et al demonstrated with chromatin conformation capture (3C) that a breast cancer risk-associated enhancer physically interacts with CITED4^8^, which is an epigenetic factor that inhibits hypoxia-activated transcription in cancer cells^21^. Similar observations have been made that connect polymorphisms at another breast cancer risk-associated enhancer to the expression of NRBF2 ^22^, and likewise for enhancers regulating MYC expression ^23^.

The assignment of variant-associated enhancers to target genes is often not obvious. Sophisticated computational and experimental strategies have been developed to link enhancers to their direct (or primary) target genes ^24–26^, with the assumption that these genes directly cause the cellular phenotypes associated with variants. However, indirect (or secondary) targets can also play an important role. For example, a risk-associated enhancer for persistent fetal hemoglobin (HbF) alters the expression of the transcriptional repressor BCL11A ^27^. However, it is not BLC11A *per se* that is directly responsible for the phenotype. Rather, it is de-repression of the BCL11A target gene HbF that elicits the phenotype ^28^. Highlighting an example more relevant to breast cancer, non-coding somatic mutations of enhancers alter the expression of ESR1 and downstream pathways in breast cancer ^29^. ESR1 encodes estrogen receptor (ER), a ligand-responsive transcription factor that plays an important role in cell proliferation, survival and metastasis ^30^. Outside of breast cancer, non-coding somatic mutations of enhancers in ovarian cancer regulate the expression of the transcription factors ZSCAN16 and ZSCAN12 ^31^. In summary, enhancers can indirectly influence downstream genes and pathways that drive cancer development. Since these indirect interactions are difficult to predict computationally, functionally assessing the target genes of variant-associated enhancers by direct experimental perturbation coupled to a transcriptome-wide readout is an alternative strategy. We and others have recently applied single-cell CRISPRi screens with a transcriptomic readout (Perturb-seq) to directly measure the primary and secondary target genes of enhancers ^32–37^. These analyses show that, through the action of secondary gene targets, enhancers can have far-reaching effects on gene regulatory networks and these effects can be used to interpret enhancer variants. Together, these data suggest that identifying primary and secondary targets can elucidate the roles of variant-associated enhancers in cancer.

In this study, we seek to investigate the function of enhancers which are associated with breast cancer variants. Using single-cell CRISPRi screens, we identified variant-containing enhancers that directly and indirectly regulate cancer genes and programs. By revealing networks of enhancer connections with their indirect target genes, we identify enhancer network motifs with relevance to breast cancer. Our study demonstrates that enhancers associated with breast cancer form regulatory networks that link cancer genes across the entire genome, providing a more comprehensive understanding of the contribution of enhancers to breast cancer development.

## RESULTS

### Single-cell CRISPRi screens of breast cancer associated regulatory elements

To functionally characterize the enhancers associated with breast cancer risk (**Fig. 1a**), we prioritized perturbation of regulatory elements linked to Genome-Wide Association Study (GWAS) variants and noncoding somatic mutations (**Fig. 1b**). First, a meta-analysis of breast cancer GWAS datasets from diverse populations identified 4,452 variants (SNPs, single nucleotide polymorphism) in 142 distinct loci^8^. In order to identify active regulatory elements, we profiled chromatin accessibility with ATAC-seq in 8 breast cancer cell lines (**Fig. 1b**) and integrated this with publicly available H3K27ac ChIP-seq datasets^17^ (**Supplementary Table 1**). Second, the Pan-Cancer Analysis of Whole Genomes (PCAWG) Consortium sequenced 123 breast cancer patients to identify 787,212 non-coding somatic mutations across the human genome^10^. We prioritized regulatory elements that may drive cancer as those that contain variants from two or more patients and are enriched for open chromatin and H3K27ac enhancer features (**Fig. 1b-d**, see Methods). In total, these analyses identified 3,512 regulatory elements for perturbation (2,123 putative enhancers, 1,389 promoters) (**Supplementary Table 2**). GWAS-associated perturbed regions do not necessarily harbor variants, but all regions selected in the somatic mutation screen have patient-derived variants (**Supplementary Fig. 1a-b**). Furthermore, the ATAC-seq signals and H3K27ac ChIP-seq signals were quantified for each perturbed region (Fig. 1b, **Supplementary Fig. 1c-d**). The 3,512 non-coding regulatory elements are distributed throughout the entire genome (**Supplementary Fig. 1e**).

**Figure 1:**
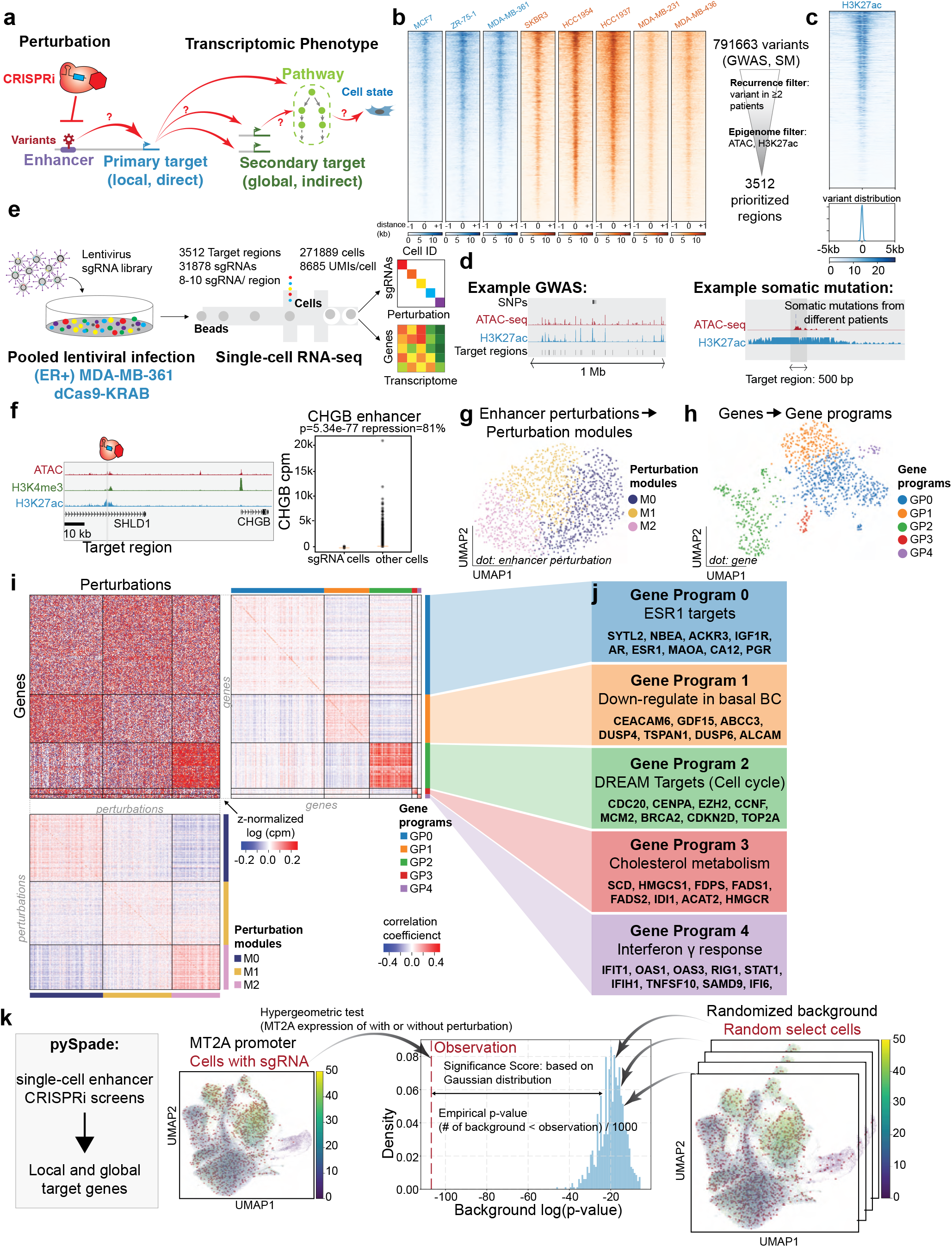
Genome-wide CRISPRi screens of breast cancer associated enhancers. a. Schematic overview of this study using single-cell CRISPRi screen to connect variant-associated enhancers with downstream gene targets and pathways. b. (Left) ATAC-seq signals of all 8 breast cancer cell lines across 3,512 perturbation regions. Cell lines colored in blue indicate the subtype of ER positive, and cell lines in orange indicate the subtype of ER negative. (Right) Schematic illustration of filtering the non-coding regulatory elements with breast cancer variants and epigenetic markers. c. (Upper) The H3K27ac signals of (ER+) MDA-MB-361 cells across 3,512 perturbation regions. (Lower) The localization of somatic mutations in the perturbation regions. d. Genome browser shots of perturbation regions prioritized by GWAS and somatic mutations (SM). e. Single-cell CRISPRi screen overview and key statistics. sgRNA libraries were packaged into lentivirus to infect the breast cancer cells. We sequenced 558,137 cells from (ER+) MDA-MB-361 and (ER-) MDA-MB-231 cell lines and generated single-cell transcriptome and perturbation data. f. sgRNAs targeting CHGB enhancers demonstrate 81% of repression in (ER+) MDA-MB-361 cells, indicating robust CRISPRi in (ER+) MDA-MB-361 cells. (Raw p-value, Student’s t-test). (Box plot: center line, median; box limits, top and bottom 10%; whiskers, 1.5x interquartile range; points, outlier). g. The UMAP plot of somatic mutation perturbation modules in (ER+) MDA-MB-361. h. The UMAP plot of gene programs. i. (Upper left) The normalized expression heatmap arranged by perturbation modules and gene programs. (Upper right) Correlation coefficient heatmap of gene programs indicates high similarity within the same gene program. (Lower) Correlation coefficient heatmap of perturbation modules confirms that enhancers within the same module share a similar expression pattern. j. Functional annotation of each gene program identified by GSEA. k. Schematic overview of the pySpade package to identify direct and indirect gene targets from CRISPRi-mediated repression of enhancers. To reduce false positives, we modeled background distributions by random simulation. The figure illustrates an example of how MT2A promoter perturbation impacts MT2A expression. We compare the MT2A expression level of cells with MT2A promoter sgRNAs to all the other cells by hypergeometric test. We implement a background correction strategy, that randomly selects cells 1,000 times from the population and calculates the p-value of MT2A expression level comparing random selected cells and other cells. Based on the background distribution and observation p-value, we calculate the empirical p-value and adjusted p-value.

We engineered stably expressing dCas9-KRAB breast cancer cell lines and tested repression efficiency (**Supplementary Fig. 2a**). Next, we performed pilot single-cell CRISPRi screens with the transcriptome as a phenotypic readout of enhancer activity (**Fig 1e**). Two important parameters for single-cell screens are 1) CRISPRi efficiency and 2) lentiviral infectivity (to maximize the number of sgRNAs per cell). We tested five breast cancer cell lines for these two parameters with shallow sequencing (**Supplementary Fig. 2b-c**), and selected the estrogen receptor positive (ER+) MDA-MB-361 cell line and estrogen receptor negative (ER-) MDA-MB-231 cell line for further analysis. We confirmed CRISPRi knock-down efficiency for the two cell lines and we captured on average 20 sgRNA and 18 sgRNA per cell, respectively (**Supplementary Fig. 2b**). We performed single-cell CRISPRi screens targeting 3,512 regulatory elements in two diverse breast cancer cell lines. This dataset spans 31,878 sgRNAs (8-10 sgRNAs per regulatory element) in 558,137 cells sequenced at high depth (average 9,081 Unique Molecular Identifiers (UMIs)/cell). We applied extensive filtering of these cells based on other quality metrics described in Methods. After all quality control filters, 383,513 cells remained for downstream analysis (**Supplementary Table 3**). Overall, we sequenced on average 1,063 cells for each perturbed region in each breast cancer cell line. These substantial datasets provide sufficient statistical power to effectively identify transcriptional phenotypes (**Supplementary Fig. 2d-e**).

We first focused on the (ER+) MDA-MB-361 cells. Supporting robust CRISPRi efficiency, we observed up to 80% repression of targeted regulatory regions (p=5.34E-77, t-test) (**Fig. 1f, Supplementary Fig. 2**). In a previous study, we observed that clonality of sequenced cells could negatively impact interpretation^38^. We confirmed that clonality is not an issue in the datasets for both cell lines (**Supplementary Fig. 3**).

### Enhancer perturbations alter shared gene programs

Single-cell screens link genetic perturbations to genome-wide transcriptional phenotypes. Previous studies ^39,40^ have identified shared gene programs that are commonly altered by multiple perturbations. If multiple distinct perturbations (defining a perturbation module) affect a common set of genes (defining a gene program), the rationale is that these perturbed genes/enhancers may contribute to breast cancer through shared mechanisms. We wondered if perturbations of variant-associated enhancers would also lead to alterations in cancer associated gene programs.

First, to identify perturbation modules that yield similar transcriptional phenotypes, we combined cells with a given regulatory element perturbed as a ‘meta-cell’ and clustered meta-cells by expression similarity ^40^. We first filtered out lowly expressed genes and focused on the top 1,000 highly variable genes. Expectedly, perturbation of enhancers does not dramatically alter global transcriptional state and we identified 3 perturbation modules with subtle transcriptional changes (**Fig. 1g**). Second, to unbiasedly identify co-regulated gene programs, we transposed the meta-cell matrix and clustered genes based on response similarity upon perturbation. We identified 5 distinct gene programs (**Fig. 1h**). Co-clustering reveals relationships between perturbation modules and gene programs (**Fig. 1i**), suggesting that distinct perturbations of enhancers lead to alterations in gene programs. Correlation analysis confirms that genes in the same gene program have similar co-expression patterns across all perturbations (**Fig. 1i**, right and bottom). We performed functional annotation with GSEA (Gene set enrichment analysis) which allows us to link gene programs to cancer-relevant pathways (**Fig. 1j**, **Supplementary Table 5**). For example, Gene Program 2 is significantly enriched for targets of the DREAM complex (p = 9.24E-272, FISCHER_DREAM_TARGETS), which regulates cell cycle genes including CDC20, CENPA, MCM2, CDKN2D, and TOP2A^41^. Furthermore, our analysis nominates regulatory elements in Perturbation Module M2 as most associated with up-regulation of Gene Program 2 (cell cycle). We observed similar patterns in both GWAS and somatic mutation screens (**Supplementary Fig. 4**). Overall, these analyses indicate that perturbation of variant-associated enhancers leads to altered expression of cancer gene programs, nominating potential dysregulated pathways for cancer development.

### Variant-associated enhancers directly regulate cancer genes in ER+ cells

To pinpoint specific genes impacted by enhancer perturbation from single-cell CRISPRi screens, we developed a computational tool, pySpade. We considered two types of targets: direct (or local/primary) target genes that are in close physical proximity (+/- 2Mb) to a perturbed region, and indirect (or global/secondary) target genes are all other genes with altered expression (**Fig. 1k**, Supplementary Fig. 5a). For example, consider CRISPRi perturbations of the MT2A promoter. We compared MT2A expression in cells with sgRNAs targeting MT2A promoter against all the other cells without MT2A promoter perturbation. To reduce false positive hits, we implemented a background correction strategy that uses random sampling to account for biases associated with different genes, targeted regions, and cell states (**Supplementary Fig. 5b-c**). Cluster analysis indicates that the number of sgRNAs detected in a cell varies by transcriptional state. For example, cells with high mitochondrial content tend to have fewer sgRNAs recovered. Thus, to test if a foreground of sgRNA-perturbed cells exhibits differential gene expression, it should be compared to an appropriate background of cells that also contain sgRNAs. To account for this, our background sampling strategy randomly selects cells weighted by sgRNA content. We use this information to calculate a Significance Score that encapsulates differences between observed changes in gene expression and random changes. We compared multiple methods to estimate significance, and chose a Gaussian approximation for its dynamic range and low computational cost (see Methods).

In (ER+) MDA-MB-361 cells, pySpade identified 369 non-coding regulatory regions with primary target genes (Significance Score < -15; fold change > 25%) (**Fig. 2a-b, Supplementary Table 6**). Consistent with the function of enhancers and promoters, we observe that CRISPRi-mediated repression of enhancers leads to downregulation of target gene expression (**Fig. 2a, volcano plot**). Expectedly, CRISPRi perturbation of the promoters of breast cancer associated genes leads to their down-regulation. 84 out of 369 local gene hits have promoters within 2 kb of CRISPRi target regions. These include MLLT10, BRCA2 and MUC1, which are cancer genes in the COSMIC database (**Supplementary Fig. 6a-f**). In addition, we also found 40 local gene hits and their perturbation regions are located within the same contact domain of another ER+ breast cancer cell line MCF7 ^42^. Several hits also coincide with previously reported eQTLs in breast cancer patients, highlighting the relevance of our in vitro model ^43–45^ (**Supplementary Table 6**). Next, we examined instances where variant-associated enhancers have primary targets with literature-supported links to cancer.

**Figure 2:**
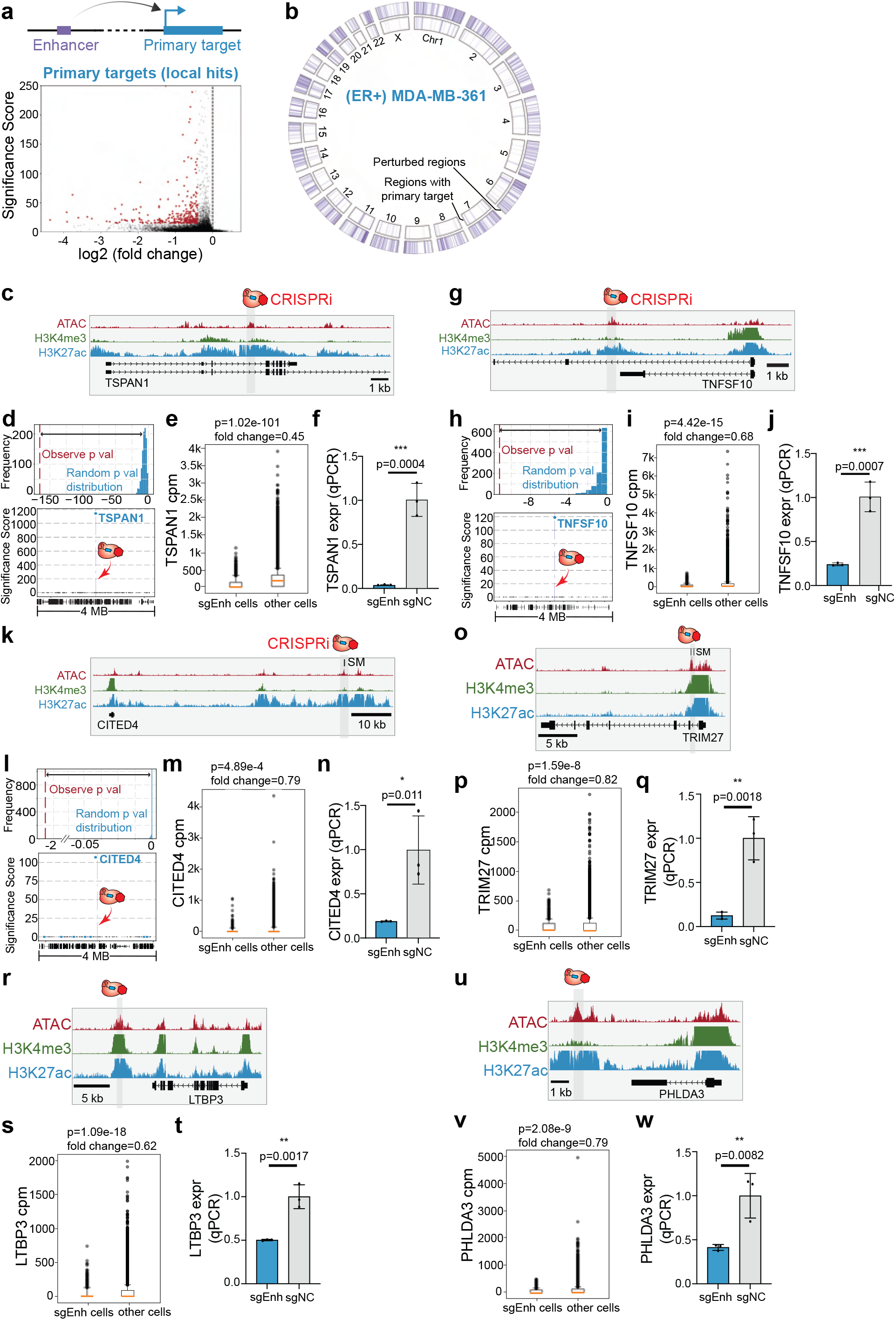
Variant-associated enhancers directly regulate cancer genes in ER+ cells. a. Enhancers can directly regulate the primary target genes. The volcano plot shows 366 primary targets identified in (ER+) MDA-MB-361 cells. b. Circos plot illustrating all perturbed regions (3,512) and regions with local hits (366). c. Genome Browser snapshot of the TSPAN1 locus, with targeted enhancer indicated. d. (Top) The randomized background distribution and the observed p-value of TSPAN1. (Bottom) A Manhattan plot of differential expression p-values +/- 2-Mb from the perturbed enhancer shows TSPAN1 is the only significant gene upon perturbation. e. Single-cell data shows that cells with sgRNAs targeting TSPAN1 enhancer lose 55% of TSPAN1 expression. (Raw p-value, Student’s t-test). (Box plot: center line, median; box limits, top and bottom 10%; whiskers, 1.5x interquartile range; points, outlier). f. Bulk qPCR validation shows that enhancer knock-down significantly reduces TSPAN1 expression. (NC: non-target control sgRNA, 3 biological replicates, * p< 0.05, ** p< 0.01, *** p< 0.001, Student’s t-test). g. Genome Browser snapshot of the TNFSF10 locus, with targeted enhancer indicated. h. (Top) The randomized background distribution and the observed p-value of TNFSF10. (Bottom) A Manhattan plot of differential expression p-values +/- 2-Mb from the perturbed enhancer shows TNFSF10 is the only significant gene upon perturbation. i. Single-cell data shows that cells with sgRNAs targeting TNFSF10 enhancer lose 32% of TNFSF10 expression. (Raw p-value, Student’s t-test). (Box plot: center line, median; box limits, top and bottom 10%; whiskers, 1.5x interquartile range; points, outlier). j. Bulk qPCR validation shows that enhancer knock-down significantly reduces TNFSF10 expression. (NC: non-target control sgRNA, 3 biological replicates, * p< 0.05, ** p< 0.01, *** p< 0.001, Student’s t-test). k. Genome Browser snapshot of the CITED4 locus, with targeted enhancer indicated. l. (Top) The randomized background distribution and the observed p-value of CITED4. (Bottom) A Manhattan plot of differential expression p-values +/- 2-Mb from the perturbed enhancer shows CITED4 is the only significant gene upon perturbation. m. Single-cell data shows that cells with sgRNAs targeting CITED4 enhancer lose 21% of TNFSF10 expression. (Raw p-value, Student’s t-test). (Box plot: center line, median; box limits, top and bottom 10%; whiskers, 1.5x interquartile range; points, outlier). n. Bulk qPCR validation shows that enhancer knock-down significantly reduces CITED4 expression. (NC: non-target control sgRNA, 3 biological replicates, * p< 0.05, ** p< 0.01, *** p< 0.001, Student’s t-test). o. Genome Browser snapshot of TRIM27 enhancer. p. Single-cell data comparing cells with TRIM27 enhancer and all the other cells shows TRIM27 loss of expression upon TRIM27 enhancer perturbation. (Raw p-value, Student’s t-test). (Box plot: center line, median; box limits, top and bottom 10%; whiskers, 1.5x interquartile range; points, outlier). q. The bulk qPCR validation of TRIM27 enhancer. (NC: non-target control sgRNA, 3 biological replicates, * p< 0.05, ** p< 0.01, *** p< 0.001, Student’s t-test). r. Genome Browser snapshot of LTBP3 enhancer. s. Single-cell data of LTBP3 expression comparing LTBP3 enhancer cells and the other cells. (Raw p-value, Student’s t-test). (Box plot: center line, median; box limits, top and bottom 10%; whiskers, 1.5x interquartile range; points, outlier). t. The bulk qPCR validation of LTBP3 enhancer. (NC: non-target control sgRNA, 3 biological replicates, * p< 0.05, ** p< 0.01, *** p< 0.001, Student’s t-test). u. Genome Browser snapshot of PHLDA3 enhancer. v. Single-cell data of PHLDA3 expression in PHLDA3 enhancer perturbation cells and all the other cells. (Raw p-value, Student’s t-test). (Box plot: center line, median; box limits, top and bottom 10%; whiskers, 1.5x interquartile range; points, outlier). w. The bulk qPCR validation of PHLDA3 enhancer. (NC: non-target control sgRNA, 3 biological replicates, * p< 0.05, ** p< 0.01, *** p< 0.001, Student’s t-test).

For example, TSPAN1 can promote breast cancer growth by modulating the PI3K/AKT pathway^46^. TSPAN1 also plays a role in migration and invasion in prostate cancer^47^ and gastric cancer^48^. Our single-cell screen identified an enhancer ∼10-kb downstream of the TSPAN1 transcription start site (TSS) that directly regulates its expression^46^ (**Fig. 2c**). Highlighting the specificity of this enhancer, TSPAN1 is the only differentially expressed gene within +/- 2-Mb of the perturbed region (**Fig. 2d**). Cells with sgRNAs targeting the TSPAN1 enhancer exhibit a significant 55% loss of TSPAN1 expression (p=1.02E-101, t-test) (**Fig. 2e**). To independently confirm this result, we performed bulk CRISPRi validation experiments and observed significant TSPAN1 repression upon perturbing TSPAN1 enhancer (96% repression, p=0.0004, t-test) (**Fig. 2f**).

Another cancer gene example is TNFSF10, which encodes TNF-related apoptosis-inducing ligand (TRAIL) protein. TRAIL is able to induce cancer-specific apoptosis ^49–52^, making it a potential therapeutic target for breast cancer treatment^52^. We identified an enhancer 8-kb downstream of the TNFSF10 promoter that regulates its expression (**Fig. 2g**). Single-cell analysis shows that the perturbed enhancer specifically regulates TNFSF10 expression (**Fig. 2h**), and cells with sgRNAs targeting the enhancer have a significantly lower expression of TNFSF10 (32% repression, p=4.42E-15, t-test) (**Fig. 2i**). We further validated with bulk CRISPRi and confirmed that TNFSF10 loses its expression with TNFSF10 enhancer perturbation (76% repression, p=0.0007, t-test) (**Fig. 2j**).

CITED4 is a transcriptional co-activator which can bind to CREB binding protein (CBP) and p300 to regulate gene expression^53^. CITED4 also has multiple roles in cancer development, including the modulation of cell proliferation in colorectal cancer ^54^. CITED4 also regulates the hypoxia pathway in breast cancer^21^, which is associated with cancer migration and angiogenesis. In addition, CITED4 is a strong dependency in (ER+) MDA-MB-361 cells reported by the DepMap ^55^. The single-cell screen identified an enhancer ∼60-kb upstream of the CITED4 TSS that specifically controls CITED4 expression (**Fig. 2k-l**). In addition, cells with CITED4 enhancer perturbation show significant reduction of CITED4 expression compared to control cells (21% repression, p=4.89E-4, t-test) (**Fig. 2m**). We confirmed with qPCR that CITED4 is repressed with this enhancer perturbation (81% repression, p=0.011, t-test) (**Fig. 2n**).

Several additional examples also highlight the identification of enhancers relevant to breast cancer. For example, TRIM27 is reported in the COSMIC database ^56–58^ as an oncogene in multiple types of cancer. TRIM27 can activate the epithelial-mesenchymal transition process in colorectal cancer ^59^ and regulate proliferation in ovarian cancer cells ^60^ through the AKT signaling pathway. In breast cancer, TRIM27 can repress cell senescence to promote cancer development^61^. We identified an enhancer which is close to the TRIM27 promoter (2.5 kb) that can regulate its expression (**Fig. 2o**). Local analysis confirmed the specificity of this enhancer and single-cell data shows TRIM27 repression upon enhancer perturbation (**Fig. 2P**). Loss of TRIM27 expression with enhancer perturbation is also validated with bulk assay (88% repression, p=0.0018, t-test) (**Fig. 2q**). We also observed and validated a similar phenotype with LTBP3 and PHLDA3 enhancer perturbation. LTBP3 is a cancer gene involved in the early cancer metastasis process ^62^ (50% repression, p=0.0017, t-test) (**Fig. 2r-t**). PHLDA3 is able to regulate the AKT pathway in cancer cells ^63,64^ (59% repression, p=0.0082, t-test) (**Fig. 2u-w**). Although the fold change of TRIM27 enhancer (18%) and PHLDA3 enhancer (19%) do not meet the cutoff for the screen, we are still able to validate by qPCR, suggesting weaker perturbations may also be hits. **Supplementary Table 6** provides both unfiltered and filtered differential expression analysis results.

### Variant-associated enhancers indirectly regulate secondary target genes in ER+ cells

While enhancers act directly as cis regulators, downstream effects can indirectly impact expression in trans. For example, an enhancer of the transcription factor BCL11A indirectly regulates the expression of fetal hemoglobin genes ^65,66^, and an enhancer harboring breast cancer associated variants regulates ESR1 and further regulates cancer regulated pathways^29^. In this way, variant-associated enhancers can extend their reach outside of their local genomic context. By using whole transcriptome data, we can evaluate the global effects of enhancer perturbation. Our analysis identified 9,404 enhancer-gene pairs that are putative global targets in (ER+) MDA-MB-361 cells (Significance Score < -50; fold change > 20%; expressed in > 5% cells) (**Fig. 3a, Supplementary Table 6-7**). For the putative local hits and global hits, the majority of the sgRNAs targeting the same perturbed region are functionally consistent (**Supplementary Fig. 7**). Globally, the direct and indirect connections between enhancers and target genes define an enhancer regulatory network (**Fig. 3b**). We found that the network spans all chromosomes, and 7% of genes (3,851) are differentially expressed across all perturbations. Most of the perturbations have few connections to genes (median=1); however, some genes are highly connected by different perturbations like IFI6 (**Supplementary Fig. 8a-b**). We observed that indirect targets are both up- and down-regulated across the genome, suggesting a variety of regulatory mechanisms. To explore these interactions in the context of cancer, we next examined examples of variant-associated enhancers that have cancer genes as indirect targets.

**Figure 3:**
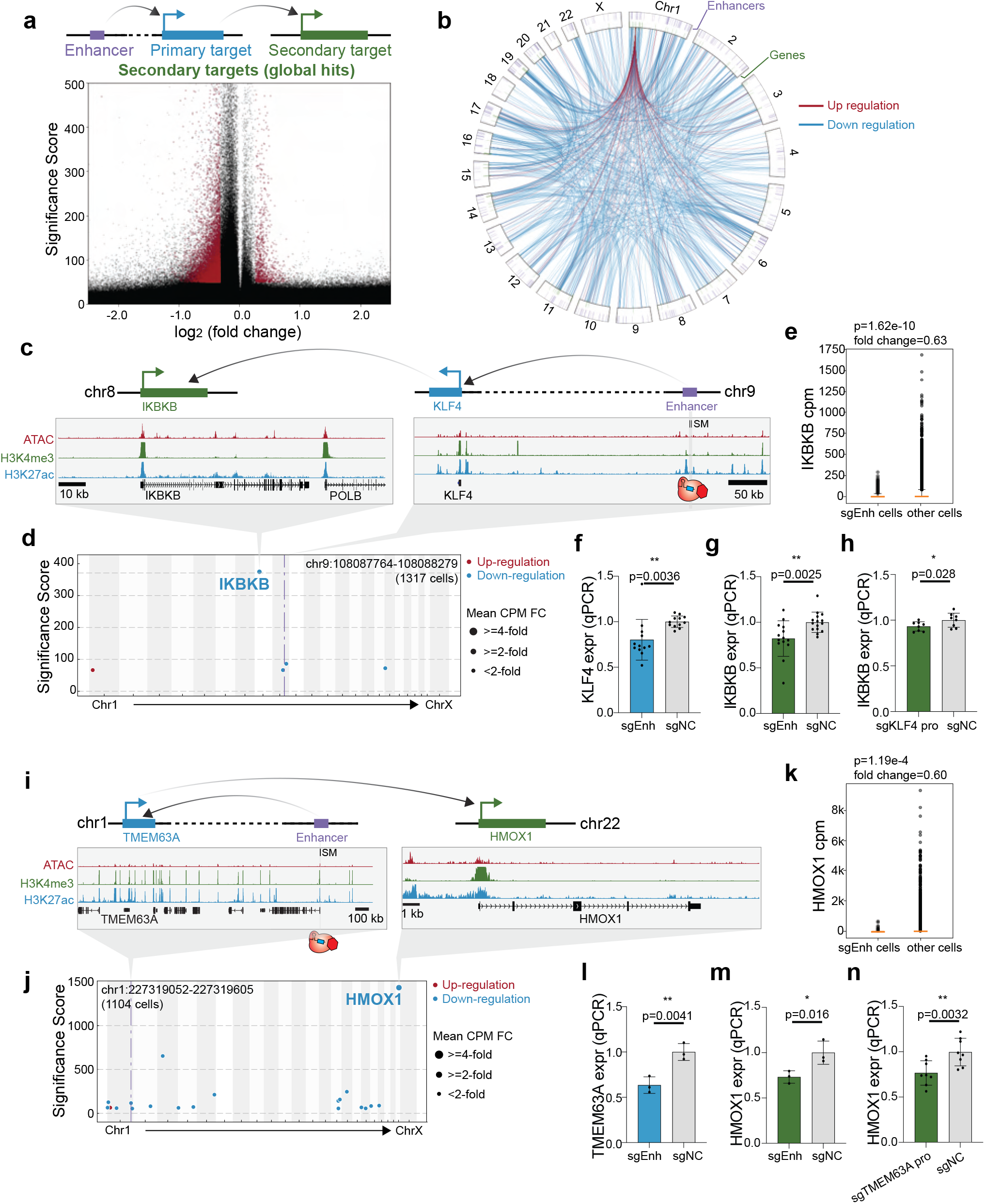
Variant-associated enhancers indirectly regulate secondary target genes in ER+ cells. a. (Top) Schematic illustration of the direct (primary) and indirect (secondary) gene targets of an enhancer. (Bottom) The volcano plot shows that most indirect gene targets are down-regulated. b. Circos plot illustrating direct and indirect gene targets of perturbed enhancers. The p-value cutoff (<10^-100^) and fold-change cutoff (> 25%) are applied to global hits. The outer ring shows the location of perturbed enhancers, the inner ring indicates the genes and the lines inside the circle show the direction of enhancer regulation. c. Genome Browser snapshots of the KLF4 (chr9) and IKBKB (chr10) loci, with targeted enhancer indicated. d. Manhattan plot of KLF4 enhancer perturbation shows that IKBKB is the most significant global hit in single-cell dataset. The dotted line indicates the perturbed region. e. Single-cell data shows that cells with sgRNAs targeting the KLF4 enhancer lose 37% of IKBKB expression compared to all the other cells. (Raw p-value, Student’s t-test). (Box plot: center line, median; box limits, top and bottom 10%; whiskers, 1.5x interquartile range; points, outlier). f. Bulk qPCR validation shows that enhancer knock-down significantly reduces KLF4 expression (direct target). (NC: non-target control sgRNA, 12 biological replicates, * p< 0.05, ** p< 0.01, *** p< 0.001, Student’s t-test). g. Bulk qPCR validation shows that KLF4 enhancer knock-down significantly reduces IKBKB expression (indirect target). (NC: non-target control sgRNA, 14 biological replicates, * p< 0.05, ** p< 0.01, *** p< 0.001, Student’s t-test). h. Knock-down primary target KLF4 promoter down-regulates secondary target IKBKB in bulk qPCR. (NC: non-target control sgRNA, 8 biological replicates, * p< 0.05, ** p< 0.01, *** p< 0.001, Student’s t-test). i. Genome Browser snapshots of the TMEM63A (chr1) and HMOX1 (chr22) loci, with targeted enhancer indicated. j. Manhattan plot of TMEM63A enhancer perturbation shows that HMOX1 is the most significant global hit in single-cell dataset. The dotted line indicates the perturbed region. k. Single-cell data shows that cells with sgRNAs targeting the TMEM63A enhancer lose 40% of HMOX1 expression compared to all the other cells. (Raw p-value, Student’s t-test). (Box plot: center line, median; box limits, top and bottom 10%; whiskers, 1.5x interquartile range; points, outlier). l. Bulk qPCR validation shows that enhancer knock-down significantly reduces TMEM63A expression (direct target). (NC: non-target control sgRNA, 3 biological replicates, * p< 0.05, ** p< 0.01, *** p< 0.001, Student’s t-test). j. m. Bulk qPCR validation shows that TMEM63A enhancer knock-down significantly reduces HMOX1 expression (indirect target). (NC: non-target control sgRNA, 3 biological replicates, * p< 0.05, ** p< 0.01, *** p< 0.001, Student’s t-test). k. n. Knock-down TMEM63A promoter down-regulates HMOX1 expression in bulk qPCR validation experiment. (NC: non-target control sgRNA, 8 biological replicates, * p< 0.05, ** p< 0.01, *** p< 0.001, Student’s t-test).

First, we identified an enhancer on chromosome 9 that indirectly regulates IKBKB on chromosome 8 (**Fig. 3c**). IKBKB encodes the kinase IKK-ß that regulates the NF-κB pathway ^67^, which has roles in proliferation, migration and apoptosis ^68,69,70^. Our single-cell transcriptome analysis revealed that perturbation of this enhancer leads to significant down-regulation of IKBKB (37% repression, p=1.62E-10, t-test) (**Fig. 3d-e**). However, we did not identify any primary target of this enhancer, likely due to the low sensitivity of single-cell analysis (**Supplementary Fig. 8c**). Thus, we used qPCR to test candidate primary target genes in the enhancer’s proximity, and identified KLF4 as putative primary targets (30% repression, p=7.18E-5, t-test) (**Fig. 3f**). KLF4 is a transcription factor, which is located ∼600-kb the enhancer, respectively (**Fig. 3c**). Importantly, qPCR experiments verified the decreased expression of global hit IKBKB when perturbing the enhancer (26% repression, p=0.0025, t-test) and the primary target KLF4 promoter (7% repression, p=0.028, t-test) (**Fig. 3g-h**).

We also identified an enhancer that indirectly regulates HMOX1, which encodes a heme oxygenase that is an essential enzyme of heme catabolism and has multiple roles in cancer development such as cell proliferation and angiogenesis ^71,72^. HMOX1 is located on chromosome 22, while the enhancer we identified is located on chromosome 1 (**Fig. 3i**). Through single-cell analysis, we found perturbation of this enhancer leads to significant loss of HMOX1 expression (40% repression, p=1.19E-4, t-test) (**Fig. 3j-k**). However, we were unable to identify local hits in the locus computationally as well (**Supplementary Fig. 8d**). qPCR analysis suggested that TMEM63A may be a direct target, which exhibited a significant loss of expression upon enhancer perturbation (37% repression, p=0.0041, t-test) (**Fig. 3l**). The protein encoded by TMEM63A has transmembrane domains and localizes in lysosomes, but the function is still unclear^73^. Moreover, we confirmed that HMOX1 exhibits loss of expression when perturbing the enhancer in bulk (27% repression, p=0.016, t-test) (**Fig. 3m**). Perturbing the primary target TMEM63A promoter also leads to secondary target HMOX1 loss of expression as well (23% repression, p=0.0032, t-test) (**Fig. 3n**). These examples support the notion that variant-associated enhancers can be indirectly connected to cancer genes throughout the genome.

### Network motifs of enhancer indirect targets in cancer cells

Since indirect targets can expand the influence of an enhancer beyond its local genomic region, we asked if these secondary interactions form network motifs. We observed one type of motif where one enhancer indirectly regulates multiple genes from the same pathway (“one to multiple”). We focused on the most significant gene program DREAM Targets and identified 19 regulatory elements across the genome that, when individually perturbed by CRISPRi, alter the expression of multiple cell cycle genes (**Fig. 4a**). As a positive control, perturbing the NFYC promoter leads to repression of cell cycle genes (**Supplementary Fig. 8e**), consistent with previous research indicating that NFYC is a known cell cycle regulator ^74,75^. Single-cell analysis shows that cell cycle genes including CHEK1 and MCM5 are significantly down-regulated upon CRISPRi of NFYC promoter (**Supplementary Fig. 8f**). These results are confirmed by bulk RNA-seq (**Supplementary Fig. 8g-h**). Another example is an enhancer that regulates cell cycle genes indirectly through PDS5B (**Fig. 4b**). The enhancer is located ∼50 kb downstream of the PDS5B promoter. PDS5B, a subunit of the cohesin complex, is essential for cohesion establishment especially for centromeric cohesion^76^. PDS5B is also essential for proper cell proliferation during development^76^. Bulk experiments confirmed that PDS5B is regulated by the enhancer (25% repression, p=0.015, t-test) (**Fig. 4c**). Global analysis of the single-cell data also confirms reduced expression of cell cycle genes including E2F6 and CCNE1 (**Fig. 4d**). As further validation of this result, bulk RNA-seq experiments also show that cell cycle genes are down-regulated after CRISPRi perturbation of the enhancer (**Fig. 4e-f**). These examples highlight a one-to-multiple relationship where one enhancer indirectly regulates multiple cancer genes.

**Figure 4:**
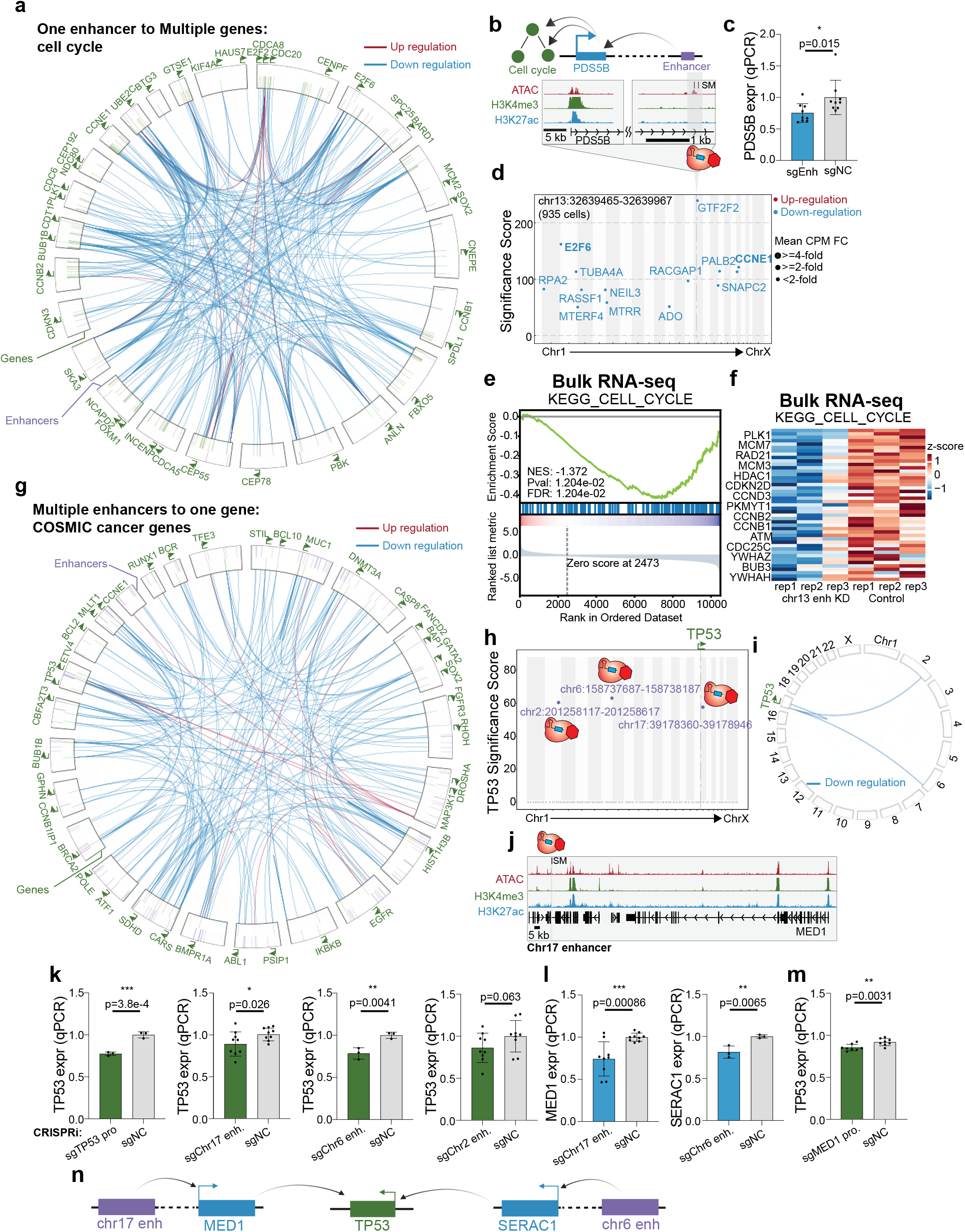
Enhancer indirect targets in cancer cells form network motifs. a. The circos plot illustrates 19 regulatory elements across the genome that, when perturbed by CRISPRi, result in altered expression of multiple cell cycle genes. These are examples of “one to multiple” enhancer motifs. Annotated genes in green belong to the cell cycle pathway. The purple lines in the outer ring indicate the localization of enhancers regulating cell cycle genes. The green lines in the inner ring indicate the target genes of the enhancers. The lines within the rings show the regulation and directionality of enhancer to gene. Knockdown of each enhancer alters the expression of multiple cell cycle genes. b. Genome browser snapshot of the PDS5B locus, with targeted enhancer indicated. This example is 1 of the 19 regulatory elements illustrated in A above. c. Bulk qPCR confirms the loss of PDS5B expression with chr13 enhancers perturbation. (NC: non-target control sgRNA, 9 biological replicates, * p< 0.05, ** p< 0.01, *** p< 0.001, Student’s t-test). d. Manhattan plot of PDS5B enhancer perturbation shows that multiple cell cycle genes lose expression. The dotted line indicates the perturbed region. e. GSEA of bulk RNA-seq validation experiment shows that the cell cycle pathway is down-regulated with PDS5B enhancer perturbation. f. The heatmap derived from bulk RNA-seq analysis shows that multiple genes of the cell cycle pathway are down-regulated with CRISPRi of the PDS5B enhancer. g. The circos plot demonstrates “multiple to one’”enhancer regulatory motifs for COSMIC cancer genes. h. The “reverse” Manhattan plot from analysis of single-cell CRISPRi screen data shows three enhancers across the genome annotated in purple that indirectly regulate TP53 expression. The y-axis indicates the adjusted p-value of TP53 in each enhancer perturbation. i. The circos plot indicates that several enhancers across the genome indirectly down-regulate TP53 in single cell data. j. The genome browser snapshot of chr17 enhancer. k. Bulk qPCR experiments confirm the indirect regulation of TP53 expression. We validated chr17 enhancer and chr6 enhancer have the ability to indirectly regulate TP53. TP53 promoter perturbation and chr6 enhancer perturbation belong to the same experimental batch and share the same sgNC control samples. (NC: non-target control sgRNA, TP53 promoter: 3 biological replicates, chr17 enhancer: 9 biological replicates, ch6 enhancer: 3 biological replicates, chr2 enhancer: 9 biological replicates, * p< 0.05, ** p< 0.01, *** p< 0.001, Student’s t-test). l. Bulk qPCR experiments identify MED1 as the primary target of chr17 enhancer and SERAC1 as the primary target of chr6 enhancer. (NC: non-target control sgRNA, chr17 enhancer: 9 biological replicates, chr6 enhancer: 3 biological replicates, * p< 0.05, ** p< 0.01, *** p< 0.001, Student’s t-test). m. MED1 promoter perturbation leads to TP53 down-regulated in bulk qPCR. (NC: non-target control sgRNA, 8 biological replicates, * p< 0.05, ** p< 0.01, *** p< 0.001, Student’s t-test). n. Schematic illustration shows that chr17 enhancer and chr6 enhancer indirectly regulate TP53 through MED1 and SERAC1 respectively.

We also observed another indirect enhancer motif where multiple regulatory elements regulate the same gene (“multiple to one”). For example, we identified cases where cancer genes defined by the COSMIC database were each regulated by multiple enhancers (**Fig. 4g**). Next, we sought to validate one example of a multiple-to-one relationship for TP53. TP53 encoding the tumor suppressor p53, which is frequently mutated in cancer ^77–79^. Single-cell analysis identifies three putative enhancers across the genome that can regulate TP53 expression indirectly (**Fig. 4h-i**). Perturbation of these enhancers lead to TP53 loss of expression with at least 30% change in the single-cell screen (**Supplementary Fig. 8i**). First, the chromosome 17 enhancer, located ∼300 kb away from its primary target MED1 (**Fig. 4j**), is able to indirectly regulate TP53, which is located on the opposite arm of chromosome 17. MED1 encodes a protein that is part of the Mediator complex and drives ER-dependent gene regulation ^80^. MED1 expression is correlated with proliferation, migration, chemoresistance ^81,82^, and poor prognosis in breast cancer patients. Second, an enhancer on chromosome 6 is able to indirectly regulate TP53 through SERAC1, which is ∼400 kb away from the enhancer (**Supplementary Fig. 8j**). SERAC1 encodes a phosphatidylglycerol remodeling protein mediating phospholipid change. Additionally, it is also a known biomarker of breast cancer ^83,84^, but the mechanisms of regulating other genes in trans remains unknown. A third enhancer is located on chromosome 2. Independent bulk qPCR experiments validate that the chromosome 17 enhancer and chromosome 6 enhancer indirectly regulate TP53 as shown in the single cell screen but not the chromosome 2 enhancer (TP53 promoter: 23% repression, p=3.8E-4, t-test. chr17 enhancer: 11% repression, p=0.026, t-test. chr6 enhancer: 22% repression, p=0.0041, t-test. chr2 enhancer: 14% repression, p=0.063, t-test.) (**Fig. 4k**). We also validate the direct regulation of these enhancers to their primary targets (chr17 enhancer: 26% repression, p=0.00086, t-test. chr6 enhancer: 19% repression, p=0.0065, t-test) (**Fig. 4l**). Furthermore, CRISPRi-mediated repression of the MED1 promoter confirms reduced expression of the secondary target TP53 (7% repression, p=0.0031, t-test) (**Fig. 4m**). These results suggest that variant-associated enhancers across the genome can alter the expression of tumor suppressors, which provides insights about how cancer genes are regulated (**Fig. 4n**).

Overall, we demonstrated that breast cancer associated enhancers are globally connected to cancer genes, which form an enhancer regulatory network.

### Identifying variant-associated enhancer networks in ER- cells

(ER+) MDA-MB-361 and (ER-) MDA-MB-231 represent two distinct subtypes of breast cancer based on estrogen receptor (ER) expression ^85,86^. Using chromatin signatures to map enhancers, previous studies demonstrated that enhancer activity is breast cancer subtype specific ^17,87,88^. On average, chromatin signatures identify 72% of putative enhancers in (ER+) MDA-MB-361 cells and (ER-) MDA-MB-231 cells as subtype specific (**Fig. 5a**). Consistent with this subtype specificity of transcriptional programs, previous studies show that both transcriptome and epigenome landscapes are distinct between ER+ and ER- patients ^89,90^, single cell RNA-seq reveals transcriptome diversity between different subtypes of breast cancer ^91^, and risk loci identified in GWAS show specificity between ER+ and ER- patients ^8^. To examine enhancer specificity in different breast cancer molecular subtypes, we repeated the single-cell CRISPRi screen targeting the same genomic regions in (ER-) MDA-MB-231 (**Fig. 5b**). Next, we assessed the global gene regulatory changes after CRISPRi perturbation of regulatory elements by repeating the analysis of perturbation modules and gene programs in MDA-MB-231 cells (**Supplementary Fig. 9**). The “DREAM Targets”program of cell cycle genes was shared between MDA-MB-361 cells and MDA-MB-231 cells. In contrast, gene programs specific to MDA-MB-231 cells include cell migration, TNFA signaling and ribosomes (**Supplementary Table 5**). These analyses suggest subtype-specific transcriptional responses upon regulatory element perturbation. We performed single cell clustering analysis and identified different cell states in MDA-MB-231 cells as well (**Supplementary Fig. 10**). By applying the pySpade analysis pipeline and adjusting with cell states, we identified 335 local hits and 9,432 putative global hits with the same filters applied to MDA-MB-361 screens (**Fig. 5c-d**). Consistent with results from (ER+) MDA-MB-361 cells, we observed that enhancer repression predominantly results in gene silencing of local and global target genes in (ER-) MDA-MB-231 cells (**Fig. 5d**). This suggests that breast cancer associated enhancers are more likely to regulate activators, in both ER+ cells and ER- cells.

**Figure 5:**
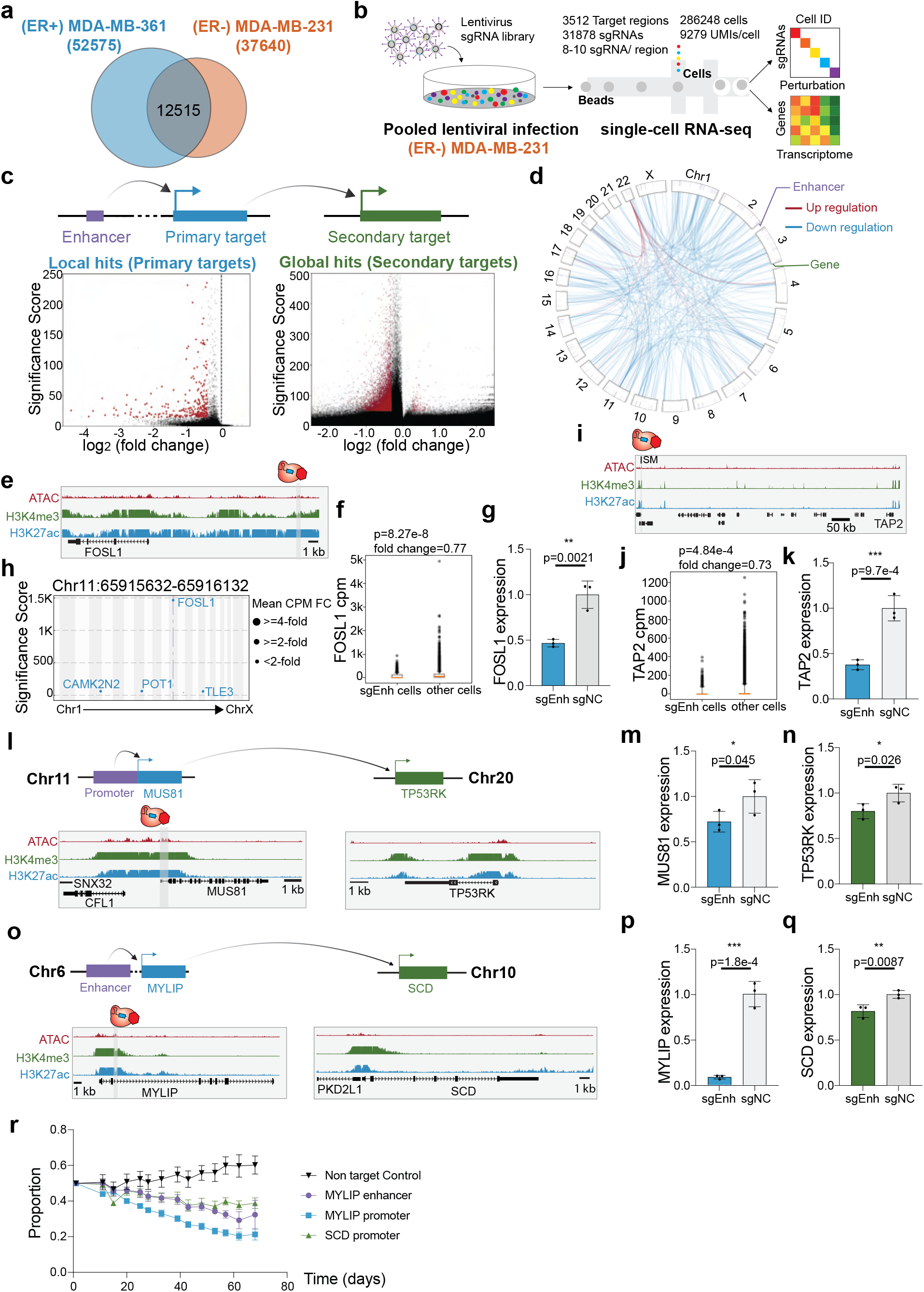
Identifying variant-associated enhancer networks in ER- cells. a. Overlap of enhancers between (ER+) MDA-MB-361 cells and (ER-) MDA-MB-231 cells. Enhancers are defined by overlapping ATAC-seq and H3K27ac ChIP-seq signals and subtracting the known transcription start site. 24% of MDA-MB-361 enhancers and 33% of MDA-MB-231 enhancers are shared. b. The workflow and statistics overview of single-cell CRISPRi screen in ER- cell line MDA-MB-231. c. Volcano plots of local hits (direct regulation) and global hits (indirect regulation). d. Direct and indirect regulation of enhancers form enhancer networks in MDA-MB-231 cells. The p-value cutoff (<10^-100^) and fold-change cutoff (>25%) are applied to global hits. e. Genome Browser snapshot of the FOSL1 locus in MDA-MB-231 cells, with targeted enhancer indicated. f. Single-cell data shows that cells with sgRNAs targeting FOSL1 enhancer lose 23% of FOSL1 expression. (Raw p-value, Student’s t-test). (Box plot: center line, median; box limits, top and bottom 10%; whiskers, 1.5x interquartile range; points, outlier). g. Bulk qPCR validation shows that enhancer knock-down significantly reduces FOSL1 expression. (NC: non-target control sgRNA, 3 biological replicates, * p< 0.05, ** p< 0.01, *** p< 0.001, Student’s t-test). h. Manhattan plot of global transcriptome change. Multiple genes are down-regulated with FOSL1 enhancer knock-down. i. Genome Browser snapshot of the TAP2 locus, with targeted enhancer indicated. j. Single-cell data shows that cells with sgRNAs targeting TAP2 enhancer lose 27% of TAP2 expression. (Raw p-value, Student’s t-test). (Box plot: center line, median; box limits, top and bottom 10%; whiskers, 1.5x interquartile range; points, outlier). k. Validation with bulk qPCR shows that TAP2 is significantly down-regulated with TAP2 enhancer perturbation. (NC: non-target control sgRNA, 3 biological replicates, * p< 0.05, ** p< 0.01, *** p< 0.001, Student’s t-test). l. Genome Browser snapshots of the MUS81 (chr11) and TP53RK (chr20) loci, with targeted enhancer indicated. m. Bulk qPCR validation shows that MUS81 is down-regulated upon CRISPRi of MUS81 promoter (direct target). (NC: non-target control sgRNA, 3 biological replicates, * p< 0.05, ** p< 0.01, *** p< 0.001, Student’s t-test). n. Bulk qPCR validation shows MUS81 promoter knock-down reduces the TP53RK expression (indirect target). (NC: non-target control sgRNA, 3 biological replicates, * p< 0.05, ** p< 0.01, *** p< 0.001, Student’s t-test). o. Genome Browser snapshots of the MYLIP (chr6) and SCD (chr10) loci, with targeted enhancer indicated. p. Validation with bulk qPCR shows that MYLIP is significantly down-regulated with MYLIP enhancer perturbation (direct target). (NC: non-target control sgRNA, 3 biological replicates, * p< 0.05, ** p< 0.01, *** p< 0.001, Student’s t-test). q. Bulk qPCR validation shows that SCD is down-regulated with MYLIP enhancer perturbation (indirect target). (NC: non-target control sgRNA, 3 biological replicates, * p< 0.05, ** p< 0.01, *** p< 0.001, Student’s t-test). r. A flow cytometry-based proliferation assay demonstrates that MDA-MB-231 cells with MYLIP enhancer perturbation exhibit cell decreased proliferation over time.

First, we examined examples of enhancers with cancer genes as primary targets. FOSL1 forms part of the AP-1 transcription factor complex and regulates multiple cancer pathways. For example, FOSL1 enhances the drug resistance in breast cancer ^92^ and is a key regulator of proliferation in triple negative breast cancer cells ^17^. We identified an enhancer 4-kb upstream of FOSL1 (**Fig. 5e**), and we observed in single-cell analysis that CRISPRi-mediated repression of this enhancer causes robust knockdown of FOSL1 expression (**Fig. 5f**). Cells with sgRNAs targeting the FOSL1 enhancer exhibit a significant 23% loss of FOSL1 expression (p=8.27E-8, t-test). This result was confirmed with independent qPCR experiments (53% repression, p=0.002, t-test) (**Fig. 5g**). Since FOSL1 functions as a transcription factor, we next asked if global gene targets were differentially expressed as a result of enhancer CRISPRi. We identified multiple putative global gene targets that had significant differential expression (**Fig. 5h**). Notably, all of these global targets were downregulated upon enhancer CRISPRi, suggesting that FOSL1 acts as an activator of these genes in (ER-) MDA-MB-231 cells. Next, TAP2 is a member of the ABC family of transporters that plays an important role in antigen presentation, with relevance to immunotherapy and multidrug resistance in cancer ^93–95^. We identified an enhancer ∼640 kb downstream of TAP2 that specifically regulates TAP2 expression (**Fig. 5i**). In our single-cell data, perturbed cells exhibit 27% repression of TAP2 expression (p=4.84E-4, t-test) (**Fig. 5j**), which we confirmed by qPCR (62% repression, p=9.7E-4, t-test) (**Fig. 5k**).

Second, we performed analysis of indirect target genes in (ER-) MDA-MB-231 cells. We expanded the analysis to the whole transcriptome to identify enhancers that can indirectly regulate cancer dependencies defined by DepMap ^55^. TP53RK, which is located on chromosome 20, encodes a kinase that enhances TP53 activity by phosphorylating p53 at Ser 15 ^96^. We observed that upon CRISPRi-mediated repression of the MUS81 promoter on chromosome 11, TP53RK expression is notably down-regulated (**Fig. 5l**). Independent qPCR experiments validating MUS81 knockdown (28% repression, p=0.045, t-test) coincides with reduced TP53RK expression (20% repression, p=0.026, t-test) (**Fig. 5m-n**). MUS81 plays a crucial role in DNA repair and genome integrity, and dysfunction of MUS81 is linked to cancer development ^97^. While previous studies have demonstrated the cooperation between MUS81 and TP53 in cancer cells ^98^, further investigation is required to establish the causal relationship between MUS81, TP53RK and TP53. Our data provides a possible connection between these pertinent cancer-associated genes.

We also identified the DepMap dependency gene SCD as a target of indirect enhancer regulation. SCD, located on chromosome 10, is associated with cell proliferation, migration and lipid metabolism in breast cancer ^99–101^. We observed that perturbing an enhancer downstream of MYLIP on chromosome 6 leads to reduced expression of SCD (**Fig. 5o**). MYLIP, also known as IDOL, is a regulator of LDL ^102^. Independent experiments confirmed that perturbing enhancers downstream of MYLIP leads to loss of MYLIP expression (91% repression, p=1.78E-4, t-test) and SCD (19% repression, p=0.0087, t-test) (**Fig. 5p-q**). Since SCD is a cancer dependency, we next tested if enhancer silencing alters cell proliferation. As the gene expression changes observed were mild, we modified a flow cytometry-based method to sensitively quantify changes in cellular proliferation. We observed a significant decrease in cellular proliferation upon MYLIP enhancer silencing compared to the non-targeting control (**Fig. 5r**). These results indicate that variant-associated enhancers can influence cancer phenotypes by modifying the expression of indirect target genes.

### ER+ and ER- cells show distinct transcriptome signatures

Since the (ER+) MDA-MB-361 and (ER-) MDA-MB-231 cell lines tested share active enhancers as defined by chromatin signatures^17^ (**Fig. 5a**; 28% enhancers shared), we asked if variant-associated enhancers in these two cell lines regulate similar sets of genes.

To compare the enhancer activity between these cell lines, we first focused on local hits. Only 38 local hits are common across the two cell lines (10-11% of local hits in each cell line), and 34 of them are promoters (**Fig. 6a**, Supplementary Fig. 11). In short, despite sharing some active chromatin, most of the enhancers are functionally distinct in these two cell lines (**Supplementary Table 8**). To further quantify cell type specific effects of enhancers, we compared the p-values of local hits across the two breast cancer cell lines (**Fig. 6b**). Hits located in the diagonal suggest that the most functionally shared elements between the two cell lines are promoters. In contrast, enhancer function is cell-type specific. We identified one enhancer that, when perturbed in either cell line, alters TSPAN1 expression (**Fig. 6c**), which we independently confirmed with bulk qPCR experiments (MDA-MB-361: 96% repression, p=0.0004, t-test. MDA-MB-231: 98% repression, p=4.3E-6, t-test) (**Fig. 6d**).

**Figure 6:**
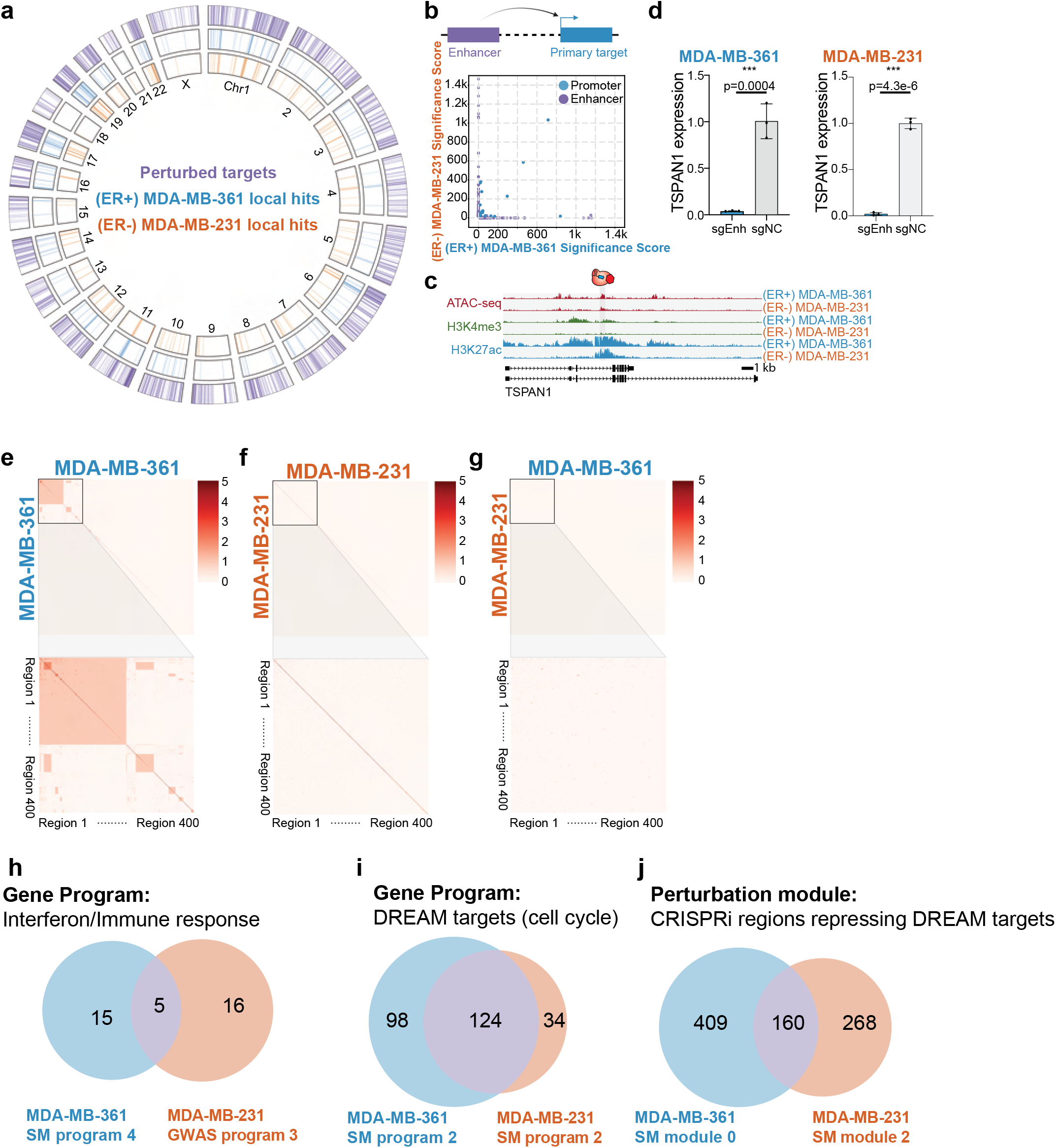
ER+ and ER- cells show distinct transcriptome signature. a. The circos plot indicates the perturbed regions and identified primary targets in (ER+) MDA-MB-361 cells and (ER-) MDA-MB-231 cells. b. The scatterplot indicates p-values of direct targets in the two cell lines. The large number of off-diagonal points indicates that enhancer expression phenotypes are significantly cell line specific. c. Genome Browser snapshots of the TSPAN1 locus in the two cell lines. d. Bulk qPCR experiments show TSPAN1 enhancer regulates TSPAN1 in both (ER+) MDA-MB-361 and (ER-) MDA-MB-231 cell lines. To facilitate comparison, we copied the MDA-MB-361 data from Figure 2F. (NC: non-target control sgRNA, 3 biological replicates, * p< 0.05, ** p< 0.01, *** p< 0.001, Student’s t-test). e. Clustering analysis of global hits between perturbed regions in (ER+) MDA-MB-361 cells. f. Clustering analysis of global hits between perturbed regions in (ER-) MDA-MB-231 cells. g. Clustering analysis of global hits between (ER+) MDA-MB-361 and (ER-) MDA-MB-231. h. The overlapping genes of MDA-MB-361 somatic mutation gene program 4 (Fig. 1j) and MDA-MB-231 GWAS gene program 3 (Supplementary Fig. 9d). Both programs are related to immune response, but only 5 genes are common in these cell lines. i. The overlapping genes of MDA-MB-361 somatic mutation gene program 2 (Fig. 1j) and MDA-MB-231 somatic mutation gene program 2 (Supplementary Fig. 9b). The DREAM target pathway is shared across two breast cancer cell lines, and many of them are overlapping. j. Comparison of perturbation modules that down-regulated DREAM targets gene program in MDA-MB-361 and MDA-MB-231 somatic mutation screens. Module 0 in MDA-MB-361 is able to down-regulate DREAM targets (Fig. 1i) while module 2 in MDA-MB-231 cells plays a similar role (Supplementary Fig. 9a). The numerical labels correspond to perturbation regions.

Next, we examined global hits across perturbations in the two cell lines. By performing pairwise analysis across perturbed regions in (ER+) MDA-MB-361 cells, we observed a cluster of regions with common global hits (**Fig. 6e**). These represent 2 highly connected interferon genes as described earlier. In (ER-) MDA-MB-231 cells, we did not observe common global hits across the perturbed regions (**Fig. 6f**). Interestingly, while interferon genes are expressed in both cell lines, they are only differentially expressed in single-cell CRISPRi screens experiments on ER+ cells, suggesting subtype-specific regulation of this pathway, perhaps mediated by ESR1^103^. Next, we repeated this analysis across (ER+) MDA-MB-361 and (ER-) MDA-MB-231 cell lines. We did not observe significant overlap of global hits when comparing the same regions in different cell lines (**Fig. 6g**). To increase the sensitivity of this analysis, we next compared perturbation modules by overlapping genes between ER+ and ER- (**Fig. 6h-i**). While immune response gene programs are enriched in both ER+ and ER- cells, only 5 genes are shared (DDX58, IFIT1, ISG15, ISG20, SAMD9) (**Fig. 6h**). In contrast, for the DREAM targets gene program, >100 genes are shared across ER+ and ER- cells. These results suggest that variant-associated regulatory regions converge to regulate the expression of common cell cycle genes in both subtypes (**Fig. 6i**). Lastly, we asked whether the same regulatory regions in ER+ and ER- are functionally similar. Focusing on perturbations that led to DREAM targets pathway down-regulation, we found that less than 50% of the regions are shared across cell lines (**Fig. 6j**). Overall, these results highlight cell line and subtype specific regulation of common cancer pathways.

## DISCUSSION

Dysregulation of diverse cellular processes can contribute to cancer development ^104,105^. Somatic mutations that alter genes are well-established drivers of this cellular dysregulation ^11,55,106,107^. For example, loss of function coding mutations can abolish the activity of tumor suppressors such as TP53, BRCA1, and BRCA2 ^108–110^. Additionally, copy number amplification of ERBB2, EGFR, or AKT2 ^111–113^ allows cancer cells to bypass growth signals and uncontrollably proliferate. By experimentally characterizing the functions of enhancers in breast cancer cells, we provide evidence that variant-associated enhancers can modify the activities of cancer genes and programs, in cis and in trans. These linkages illustrate how non-coding regions can contribute to cancer development, and highlight the need for more comprehensive mapping studies and deeper functional studies.

By measuring the transcriptional effects of systematic enhancer repression, this study yields insights into the motifs of enhancer networks in cancer (**Fig. 7**). Previous studies have linked variants to cancer development, but the intermediate molecular steps are missing. Our pipeline is able to link variants, enhancers, primary target genes, downstream genes to cancer development. We have established modes of direct enhancer regulation and three different modes of indirect enhancer regulation: one-to-one, one-to-multiple and multiple-to-one. One class of variant-associated enhancers directly activates the expression of nearby cancer genes. For example, we identified enhancers that directly regulate the cancer genes TSPAN1, CITED4, and TNFSF10. Another class of enhancers indirectly regulates the expression of cancer genes in a distal genomic locus, often on a different chromosome, by directly activating a nearby gene that has trans-regulatory activity (one-to-one). For example, we identified an enhancer of KLF4 that indirectly regulates the expression of the cancer gene IKBKB. These trans-regulatory genes include transcriptional regulators. In this way, a variant-associated regulatory element can alter the activity of multigene cellular pathways that control cell proliferation (one-to-multiple). For example, the variant-associated enhancer of PDS5B modifies the expression of multiple cell cycle genes. Interestingly, we observe instances where multiple variant-associated enhancers on different chromosomes indirectly regulate the same cancer gene (multiple-to-one). For example, enhancers on chromosome 6 and 17 indirectly regulate the expression of TP53. Overall, these observations highlight intricate regulatory networks connecting variant-associated enhancers to cancer genes across the genome (**Fig. 7**).

**Figure 7:**
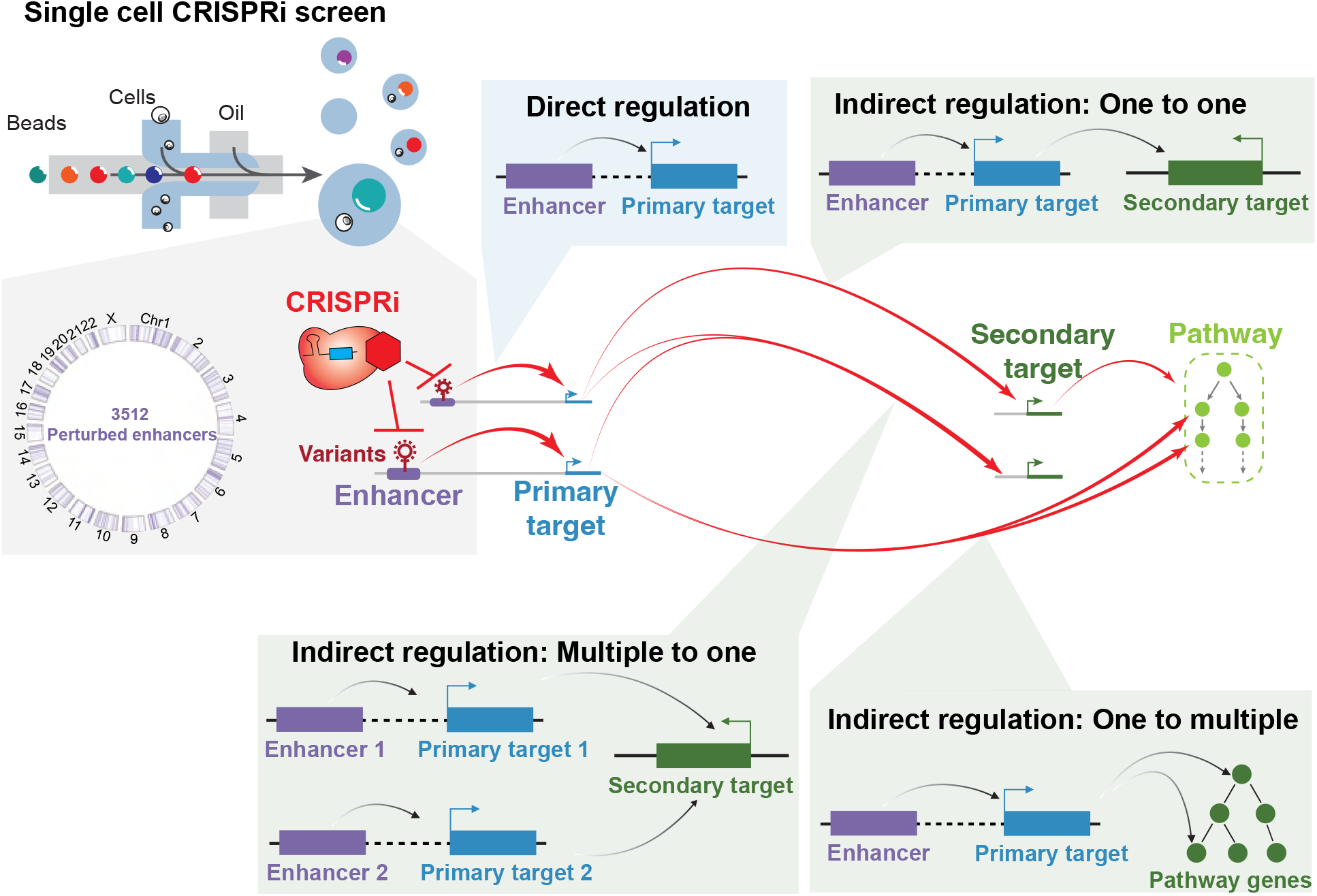
Enhancer regulatory networks connect variant-associated enhancers to global cancer genes. An overview of enhancer networks identified with Perturb-seq of breast cancer associated enhancers, including direct regulation and three types of indirect regulation.

We present pySpade, a computational pipeline for Perturb-seq differential expression analysis. pySpade globally identifies differentially expressed genes, with minimal preprocessing and optimization. It facilitates parallel and batch analysis, which will be useful as the scale of screens continues to expand. Implementing background randomization, pySpade independently accounts for the bias of each gene, target region, and cell state, which enhances the sensitivity of perturbation signal detection. However, the background randomization step can be computationally costly, and alternative approaches to model the background may be needed.

Our global analysis connects breast cancer associated enhancers to primary target genes and secondary cancer genes. Expectedly, primary target genes include genes with known roles in gene regulation. However, many primary target genes do not. For example, we identify an enhancer on chromosome 1 that indirectly alters HMOX1 expression level through its primary target TMEM63A, which has not been reported to have gene regulatory activity. These observations are consistent with other Perturb-seq studies demonstrating that perturbation of non-regulatory genes can have a significant impact on the transcriptome. For example, TMEM242 is not a known regulator, but perturbing TMEM242 leads to changes in the expression of mitochondrial pathway genes ^39^. Similarly, perturbation of the muscle contraction gene troponin T2 (TNNT2) yields a transcriptional phenotype in cardiomyocytes^114^. Thus, while Perturb-seq establishes links between cancer-associated regulatory elements and cancer genes, understanding the underlying mechanisms still requires further investigation.

As whole genome sequencing of tumors becomes increasingly common ^11,115,116^, the number of variant-associated enhancers will also rise. As most of these enhancers will have unknown links to cancer ^117^, the high throughput CRISPRi strategy used in this study is one versatile approach to functionally test variant-associated enhancers. Breast cancer development involves multiple pathways that may not be completely captured by phenotypic screens based on proliferation and invasion. In contrast to traditional CRISPR screens that use cell fitness as a phenotypic readout ^55^, one advantage of a transcriptomic readout is the ability to pinpoint the primary and secondary targets downstream of a perturbed enhancer, thus providing insights on perturbed cancer genes ^32^. However, one drawback of this CRISPRi strategy is that it lacks the resolution to give insight on variants at nucleotide resolution. Other approaches have been developed to bridge this gap and experimentally test altered sequences of potentially pathogenic enhancers. For example, saturation mutagenesis of the TERT promoter at base-level resolution in an exogenous reporter system identified specific variants that confer increased or decreased expression of TERT ^118^. Likewise, endogenous CRISPR-mediated saturation mutagenesis of the BCL11A enhancer identified the key base positions for proper regulation of fetal hemoglobin ^66,119^. Recently developed technologies including prime editing and base editing will also play important roles to address this challenge ^120,121^. Ultimately, interpreting non-coding variants will require orthogonal approaches to experimentally link enhancers and their variants to molecular and cellular phenotypes relevant to cancer.

This study motivates several future directions. First, the Perturb-seq framework can be flexibly applied to connect risk-associated enhancers to cancer genes in other cancer types and models, thereby broadening our understanding of cancer development. Second, functional analysis of patient-derived variants for the enhancers prioritized in this study can more directly connect genetic variation to breast cancer mechanisms. Third, computational modeling to predict the effects of genetic variants at these prioritized enhancers will facilitate clinical diagnosis. Fourth, genetic or chemical screens can be applied to selectively modulate the transcriptome, with the goal of converting cancer transcriptomes towards their normal counterparts. These integrated approaches will improve personalized medicine for cancer risk.

In summary, we established an integrated pipeline for the functional characterization of variant-associated enhancers and their roles in regulating breast cancer genes and cellular phenotypes. This analysis elucidates the roles of variant-associated enhancers as direct and indirect regulators of cancer genes and pathways, through diverse regulatory architectures. This resource suggests new testable hypotheses for the molecular mechanisms of enhancers and their variants in cancer development. Systematic approaches like this one will add to our understanding of the genetic drivers of cancer.

## LIMITATIONS OF THE STUDY

This study has several limitations. First, while variants identified from GWAS represent larger patient cohorts, the resolution of causal variants is low. Second, while somatic mutations identified from whole genome sequencing are high resolution, the number of patients analyzed is small, leading to incomplete catalogs of recurrently mutated enhancers. Notably, it is unclear whether the somatic mutations are causal variants. Third, this study only used two immortalized breast cancer cell lines as model systems. We expect application in more advanced models will have increased cancer relevance. More complex biological systems such as cancer organoids and xenografts that can better model the tumor microenvironment and cell-cell interactions will be needed to address this issue^105^. Fourth, we observed that CRISPRi repression of enhancers yields small changes in gene expression. This low effect size may be attributed to biological or technical factors. Biologically, enhancers only partially contribute to a gene’s full expression output. Technically, CRISPRi may not completely silence a targeted enhancer. In addition, we used a high MOI (multiplicity of infection) strategy with multiple sgRNAs per cell, which could potentially affect the sensitivity. While this issue impacts lowly expressed genes more, several solutions include deeper sequencing, sequencing more cells, and target sequencing ^122^. In addition, since protein expression is not always perfectly correlated with mRNA expression ^123^, it will be important to assess how enhancer perturbation alters protein expression and activity. Fifth, this study focuses on the impact of variant-associated enhancers by repressing whole enhancers, rather than directly examining variant function. CRISPRi is much more scalable than precision editing strategies, and gives an upper bound on enhancers that may have phenotypic consequences. It is expected that only a subset of variants alter enhancer activity, and these phenotypes will be weaker than CRISPRi-mediated phenotypes. Notably, it took two months to detect the proliferation phenotype of MYLIP enhancer knockdown. The phenotypic effect of variants may be even milder, and could take longer to develop, which has implications for the role of variants in cancer risk.

## SUPPLEMENTAL TABLE LEGENDS

**Supplementary Table 1**: The breast cancer cell lines tested in this study and their culture condition.

**Supplementary Table 2**: List of all the breast cancer lines H3K27ac and H3K4me3 quantification of each perturbation region and the sgRNAs sequences targeting those regions.

**Supplementary Table 3**: Sequencing statistics and mapping quality.

**Supplementary Table 4**: List of sgRNAs and primers for bulk validation experiments and customized library preparation primers.

**Supplementary Table 5**: List of the enriched GSEA pathways in perturbation module and gene program analysis.

**Supplementary Table 6**: Local hits and global hits of two cell lines.

**Supplementary Table 7**: List of individual sgRNA status for all the hits.

**Supplementary Table 8**: Cell line specific enhancers and target genes.

## METHODS

### Experimental details

#### Golden Gate cloning of individual sgRNA

We selected 3 sgRNAs from our screens or literature^124^ and cloned them individually for validation experiments (**Supplementary Table 4**). sgRNA oligos are synthesized by Integrated DNA Technologies (IDT). The forward strand structure is 5’-CACCG-sgRNA-3’ and the reverse strand structure is 5’-AAAC-sgRNA(reverse)-C-3’. We annealed oligos in the presence of T4 Polynucleotide Kinase (NEB). We then performed Golden Gate assembly with diluted oligos, CROPseq-Guide-Puro backbone (Addgene ID: 86708), T7 DNA ligase (NEB) and BsmbI (NEB).

#### sgRNA library construction

Oligo pools for 20,139 (GWAS screen) and 11,838 (Somatic mutation screen) sgRNAs were synthesized by GeneScript Biotech with the sequence structure: 5’-GTGGAAAGGACGAAACACCG-sgRNA-GTTTTAGAGCTAGGCCAACATGAGGATCAC-3’. The oligos were amplified using NEBNext High-Fidelity PCR master mix to make them double stranded. sgRNAs were inserted into the BsmBI-digested and gel-purified CROPseq-Guide-Puro backbone (Addgene ID: 86708) with Gibson Assembly in the ratio of 2 ug sgRNA PCR product and 3 ug digested plasmid backbone. The final plasmid library was amplified and purified through electroporation under ampicillin selection overnight. The estimated coverage of clones per sgRNA was about 1600. ZymoPURE Plasmid Gigaprep Kit was used to extract the plasmid library. We confirmed the complexity by sequencing the plasmid library and cDNA of infected cells^32^. The detailed protocol is described in Xie et al., 2019 ^125^.

#### Virus packaging, titration and infection

One million Lenti-X 293T cells were plated in 10-cm dishes one day before transfection. 16 ug of plasmid were prepared for each plate at a ratio of 4:3:1 (sgRNA library: psPAX2(Addgene ID 12259): pMD2G(Addgene ID 12260)). 48 uL of TransIT LT1 (Mirus) reagent was added to the plasmid mixture in Opti-MEM, and incubated at room temperature for 15 minutes. This mixture was gently added to cells dropwise. After 8 hours, media was removed and replaced with fresh media containing 2 mM caffeine. The medium was collected two days after transfection, filtered (0.22 um), and concentrated with Lenti-X concentrator (Takara) following the manufacturer’s manual. To maximize the infectivity, breast cancer cells were seeded in 96-well plates and infected with five different concentrations of virus with serial dilution. The highest concentration with minimum cell damage was selected for a large scale of infection. In the large scale of infection, 100,000 MDA-MB-361 cells or 50,000 MDA-MB-231 cells were seeded for one well in 24-well plates and infected with the virus on the second day. The polybrene was added in the infection step to increase infectivity. On the third day, the media was removed and added with fresh media containing blasticidin and puromycin. After two weeks of antibiotics selection, the cells were collected for experiments.

#### Cell culture

Breast cancer cell lines MCF7, ZR-75-1, MDA-MB-361, SKBR3, HCC1954, HCC1937, MDA-MB-231 and MDA-MB-436 cells were purchased from American Type Culture Collection (ATCC) and cultured with alpha modified MEM medium (Sigma) with supplement of 10% FBS, 1 mM sodium pyruvate (Gibco), 10mM HEPES (Sigma), 1X Glutamax supplement (Thermo Fisher), 1X MEM non-essential amino acid (Sigma), 1 mg/mL insulin, 1 ng/mL hydrocortisone, 25 ug/mL EGF and pen/strep at 37°C and 5% CO2 ^17^. Lenti-X 293T cells (Takara) are cultured with DMEM medium with 10% of FBS at 37°C and 5% CO2.

To repress the activity of regulatory elements, we utilized catalytic dead Cas9 (dCas9) fused with the KRAB repressor domain ^126^. To generate the stably expressed dCas9-KRAB, we packaged lenti-dCas9-KRAB-blast (Addgene ID: 89567) into lentivirus and infected all the breast cancer cell lines individually. The cells were under blasticidin selection for at least a week to form the stably expressed dCas9-KRAB lines. All cells are tested quarterly to make sure they are mycoplasma free.

#### Dual color cell proliferation assay

We generated three plasmids for the fluorescence-based cell proliferation assay. Starting with the CROPseq-Guide-Puro backbone (Addgene ID: 86708), we cloned fluorescence proteins (FP: mCherry, mNeonGreen, or BFP) to generate CROPseq-Guide(MS2)-FP-P2A-Puro. They were deposited in Addgene, and the IDs for the plasmids are 209532 (mCherry), 209533 (mNeonGreen) and 209534 (BFP).

In order to sensitively and continually quantify the proliferation rate difference of perturbations, we adapted the flow cytometry based cell proliferation method^127^. We cultured the perturbed cells and negative control (NC) cells together. Cells from each condition were tagged with different fluorescent proteins respectively. We seeded 5000 cells of each condition to one well of a six-well plate with CytoFlex SRT Sorter. Each well contains both 5,000 mNeonGreen-perturb cells and 5,000 mCherry-NC. In addition, we also perform the reciprocal experiment (5,000 mCherry-perturb and 5,000 mNeonGreen-NC) to exclude the effect of fluorescent proteins. We performed flow cytometry to quantify the proportion at each cell split, which allows us to continually quantify the perturbation effect on cell proliferation.

#### RNA extraction and RT-qPCR

Cells were collected from one well of a 6-well plate. Total RNA was extracted with Quick-RNA miniprep kit (Zymo research) followed by the manufacturer’s manual. RNA concentration was measured by Qubit RNA BR assay kit (Thermo Fisher). The same amount of input RNA for each batch was used for reverse transcription (RT) reactions with SMARTScribe™ Reverse Transcriptase (Takara) and oligo dT following the manufacturer’s manual. The expression of target genes was quantified with Power SYBR green (Thermo Fisher) and normalized with internal control ACTB. qPCR primer sequences were provided by OriGene or designed by PrimerQuest™ Tool from Integrated DNA Technologies (IDT) and checked for specificity by In-Silico PCR on the UCSC Genome Browser. All the experimental conditions contained at least 3 biological replicates, which we defined as independent virus infections. For each biological replicate, 3 technical replicates were measured in the qPCR experiments. The p-value for qPCR experiments was calculated with Student’s t-test. The blue bars in the qPCR bar charts indicate the primary target genes while the green bars indicate the secondary target genes. * p< 0.05, ** p< 0.01, *** p< 0.001.

#### Library preparation for ATAC-seq

The Omni-ATAC libraries were constructed with 50,000 cells for 8 breast cancer cell lines ^128–130^, and we prepared two replicates for each cell line. dCas9-KRAB was stably expressed in 8 breast cancer cell lines MCF7, ZR-75-1, MDA-MB-361, SKBR3, HCC1954, HCC1937, MDA-MB-231 and MDA-MB-436 cells. All cells were under blasticidin selection. In the transposition reaction, cells were incubated with 100 nM of Tn5, 1% of Digitonin and 10% of Tween-20 at 37°C for 30 minutes followed by the column cleanup with Qiagen MinElute PCR Purification Kit. We used KAPA HiFi DNA polymerase (Roche) with 7 cycles of PCR for final amplification. 1.4X SPRI beads cleanup was performed after final PCR. The ATAC-seq raw and processed data were deposited in GEO.

#### Library preparation for Single-cell CRISPRi screen

Single-cell libraries for MDA-MB-361 were built with 10x Chromium Next GEM Single-Cell 3’ HT v3.1 kits. Libraries for MDA-MB-231 were built with 10x Chromium Next GEM Single-Cell 3’ v3.1 kit. For MDA-MB-361 libraries, two million cells were prepared, stained with one set of 10x 3’ CellPlex kits, and loaded into 8 lanes of reactions based on the manufacturer’s protocol. Transcriptome and CellPlex libraries were constructed following the manufacturer’s instructions. sgRNA libraries were amplified from 50 ng of transcriptome PCR1 products using the SI primer and sgRNA enrichment primer (**Supplementary Table 4)**. Nextera indexes were added in the second round of PCR. We used 1.6X of SPRI purification to clean up the final sgRNA libraries (∼500 bp). For MDA-MB-231 libraries, cells were labeled with ten different antibodies using the cell hashing protocol (BioLegend)^131^ and loaded into 16 lanes. Transcriptome libraries were built following the 10x protocol and sgRNA libraries were constructed as described in the MDA-MB-361 library part.

#### Library preparation for bulk RNA-seq

Cells from one well of the 6-well plate were collected, and total RNA was extracted with Quick-RNA miniprep kit (Zymo research). 100 ng of RNA was used to build an RNA-seq library with the chemistry of 10x Chromium Next GEM Single-cell 3’ Reagent Kit. 10 uL of beads were used in each reaction. The remaining steps of cDNA library preparation were identical to the 10x protocol with 0.25 fold of the reaction volume.

#### Sequencing

Libraries were sequenced on the Illumina NextSeq (R1 28bp, R2 56bp, and idx1 8bp) and the Illumina NovaSeq (R1 150bp, R2 150bp, idx1 8bp). The detailed information of each library is listed in **Supplementary Table 3**.

#### Screening breast cancer cell lines for CRISPRi efficiency

We used the lentiviral strategy detailed above to generate cell lines with stably integrated dCas9-KRAB. To test CRISPRi efficiency in these lines, we used lentivirus to integrate a positive control sgRNA targeting MALAT1 (**Supplementary Fig. 2a**), with a goal of at least 50% repression. We used qPCR to test for MALAT1 expression knockdown in 7 cell lines. This analysis excluded MDA-MB-436 because of its low efficiency of repression (not meeting the 50% threshold) and SKBR3 because of its slow growth rate. We next tested the receptivity of 5 remaining cell lines for lentiviral sgRNA integration (**Supplementary Fig. 2b**). This analysis excluded MCF7 because of its low infectivity performance (<10 sgRNAs/cell). Finally, we confirmed knockdown with a single-cell RNA-Seq readout in the 4 remaining cell lines (**Supplementary Fig. 2b-c**).

### Analysis details

#### sgRNA library design

We have two enhancer selection strategies for GWAS SNPs and somatic mutations. For the GWAS sgRNA library, we clustered 4,452 breast cancer GWAS SNPs^8^ into 133 1-Mb loci genome-wide. We targeted all regulatory elements within these 133 1-Mb loci since GWAS SNPs may not be the causal variants. We identified regulatory elements by overlapping ATAC-seq peaks (generated in this study) and H3K27ac ChIP-seq signals^17^. We called the ATAC-seq peaks using the function ‘callpeak’ from macs2, and we quantified the H3K27ac ChIP-seq signals within the ATAC-seq peaks by using ‘featureCounts’. We defined active regulatory elements as ATAC-seq peaks with at least 1 log2 RPKM (Reads Per Kilobase per Million mapped reads) enrichment of H3K27ac.

For the somatic mutations screen, 787,212 somatic mutations from 123 breast cancer patients^10^ were overlapped with breast cancer ATAC-seq peaks. We focused on the open chromatin regions with recurrent somatic mutations from at least two distinct patients, which can increase the possibility of targeting causal mutations. We then applied the same filters as the GWAS screen design as described above: quantifying the H3K27ac ChIP-seq signals within ATAC-seq peaks and selecting regulatory elements with more than 1 log2 RPKM H3K27ac ChIP-seq signals.

Combining GWAS and somatic mutations, we identified 3,512 non-coding regulatory elements associated with breast cancer variants, 2,123 of them are enhancers and 1,389 of them are promoters. Next, we designed the sgRNA targeting the regulatory elements. Using blast, we reduced off-target effects by prioritizing sgRNAs with at least 3 mismatches to the genome. We aimed to identify 8-10 non-overlapping sgRNAs spanning each targeted region. Overall, there were 31,878 sgRNAs targeting 3,512 non-coding regulatory elements.

#### Single-cell RNA-Seq mapping

Single-cell transcriptome libraries were mapped to the human reference genome (hg38) with the ‘count’ function from Cell Ranger software (version 6) using the default cell number 60,000 (MDA-MB-361) and 25,000 (MDA-MB-231) based on the experimental expectation. We also mapped the CellPlex (MDA-MB-361) and Cell hashing (MDA-MB-231) libraries together with transcriptome libraries using Cell Ranger software. For sgRNA libraries, we used fba^132^ to map and filter out low UMI (Unique Molecular Identifier) sgRNA by applying a saturation curve method as described in Drop-seq^133^. We then applied the pySpade ‘process’ function to remove doublets and generated files compatible with downstream differential expression analysis.

#### Perturbation module and gene program analysis

Cells with the same perturbation are grouped together (‘meta cell’) and the average cpm for all the genes are calculated for each perturbation. Genes expressed in more than five percent of the whole population are kept for analysis. We used scanpy.pp.recipe_zheng17 to select the top 1000 highly variable genes and normalize the whole matrix.

To identify perturbation modules, we clustered the ‘meta cells’ based on their gene expression similarity. The top 50 principal components (PCs) were used for dimensionality reduction, and followed by nearest neighbor analysis (neighbor=15) and louvain clustering (resolution=0.6) to identify 3 perturbation modules. Perturbations within the same module result in similar transcriptional phenotypes. To identify these gene programs unbiasedly, the filtered and normalized ‘meta cells’ matrix from the previous step was transposed with scanpy function ‘.transcpose()’. We group the similar sets of genes together by applying the same clustering strategy: the top 50 PCs for dimensionality reduction analysis, nearest neighbor analysis (neighbor=15) and the louvain clustering method (resolution=0.4) were used to identify 5 gene programs. To further understand the biological relevance of unbiasedly identified gene programs, we performed the GSEA on the gene list of each gene program. The biological processes that showed significance for each gene program were reported in the Supplementary table 5. The notebooks and h5ad files for both cell lines for this part are deposited at the Github page for this study.

#### pySpade: overall structure

The differential expression analyses described in this manuscript are based on the outputs of pySpade, which has 4 major parts:

● process: Preprocessing of sgRNA dataset including doublets removal.
● DEobs: Performing the differential expression analysis.
● DErand: Background randomized modeling of differential expression analysis.
● local/global: identifying local (primary target) and global (secondary target) from the outputs of DEobs and DErand.

#### pySpade process: data preprocessing

This section describes the function ‘process’ of pySpade. Several input files are required for the ‘process’ function: a transcriptome matrix, a sgRNA matrix and a list of CellPlex or Cell Hashing antibody names. If there are multiple libraries of single cell RNA-seq in a screen, all the libraries are combined into one transcriptome matrix with the Cell Ranger ‘aggr’ function. The sgRNA matrix also has the same cell barcodes as the combined transcriptome matrix. This function uses multiple approaches to filter out experimental doublets. First, we remove droplets containing multiple CellPlex (or Cell Hashing) tags. We also filter out low UMI antibodies by applying the same saturation curve method as sgRNA filtering described earlier. Second, we remove droplets with an excessive amount of sgRNAs, which may be potential doublets as well. We define the sgRNA outlier number as 1.5 fold IQR (Q3 - Q1), and we exclude the cells with more than Q3 + 1.5*IQR. We save the transcriptome matrix and sgRNA matrix for all the cells passing the two filters in h5 format. The ‘process’ function also generates a statistical report about the portion of cells kept at each step, number of sgRNA per cells etc.

**pySpade DEobs/DErand: differential expression analysis and background modeling** In order to identify the differentially expressed genes of perturbation, we first calculated the raw p-value and fold change with the ‘DEobs’ function of pySpade. The transcriptome matrix is normalized to count per millions (cpm) for each cell. We use hypergeometric test (‘scipy.stats.hypergeom.sf’ function) with the parameters below to calculate the effect on gene X with region A perturbation:

x: The number of cells with gene X expression smaller than median and with sgRNAs targeting region A.
M: The number of all the cells.
n: The number of all the cells with gene X expression smaller than median.
N: The number of all the cells with sgRNAs targeting region A.

We calculate the p-values for all the genes in the genome with the parameters described above. For up-regulated genes, the log p-value is derived from scipy.stats.hypergeom.logcdf(x, M, n, N). While for the down-regulated genes the log p-value comes from scipy.stats.hypergeom.logsf(x, M, n, N). The fold change is calculated as the mean of gene X cpm in perturbation cells divided by the mean of gene X cell the other cell.

To reduce the false positive hits, we simulated the background by randomly selecting cells and performing the same hypergeometric test genome-wide (account bias for different genes) using the ‘DErand’ function of pySpade. To model the background distribution, we repeated this process 1,000 times. In order to correct the effect of sgRNA number in the cells, the probability of selecting a cell is proportional to the number of sgRNAs in the cell (account bias for cell states). Ideally, the number of cells in background modeling should be identical to the number of cells in the foreground (account bias for target regions). However, this would incur too much computational cost. Thus, to simulate the effect of cell number on each perturbation and to minimize computational cost, we ran the simulation with a binning strategy. For example, in the MDA-MB-361 dataset, we simulated between 800 and 3,100 cells in 100-cell increments. For each perturbation, we selected the simulated background from above which is closest from the cell number of that perturbation. Although we tried to minimize the computational cost, it still took a day of processing on a 256 GB node for 1,000 iterations of 1 bin in this study.

#### pySpade modeling comparison: Gaussian, Empirical and Bootstrapping

To account for the background noise, we randomly select cells with ‘DErand’ function. To adjust for background, we tried 3 methods: Gaussian, Empirical and Bootstrapping. First, we used a Gaussian distribution as an approximation to model the background hypergeometric p-values. We then compare the observed p-value with the random background to calculate an adjusted p-value. In a similar way, the adjusted fold change is calculated by dividing the cpm of observation and average cpm of all the background iterations. However, for some distributions with small standard deviation, we noticed that the adjusted p-values are likely to be inflated, and not all of them follow Gaussian distribution. The second method is empirical p-value. We calculated the proportion of background randomization p-values that are smaller than the observation p-value. For example, if one randomized background p-value is smaller than the observation p-value, the empirical p-value is 0.001 since there are 1,000 randomized p-values in the distribution. The last strategy is the bootstrapping method (**Supplementary Fig. 12a**). Since we are unable to test the distribution type of every bin and every gene, as a result, we randomly select cells with perturbation 100 times and generate an observation distribution. We then utilize KS test to characterize if the observation distribution comes from background distribution. Overall, we found that all methods can recover hits that are validated (**Supplementary Fig. 12b-g**). However, it is notable that hits called by the bootstrapping method are noisy. Empirical methods are able to recover hits, but it cannot give a good dynamic range of p-values with only 1,000 random simulations. Although both empirical and bootstrapping methods do not rely on a known statistical distribution, they require much higher computational cost. After balancing the three approaches implemented here (Gaussian approximation, empirical, and bootstrapping), we decided on the following approach: To identify the differential expressed genes, we use the Gaussian approximation since it can complete in a reasonable time scale and has reasonable dynamic range. To be statistically accurate, since the true background does not always follow a Gaussian distribution, we refer to a “Significance Score” rather than an “adjusted p-value”. In Supplementary Table 6 for the list of hits, we also include empirical p-values derived from the above analysis.

#### pySpade local/global: Identifying differentially expressed hits

The ‘local’ function focuses on the genes within +/- 2MB of perturbation, while the ‘global’ function performs the analysis with the whole genome. All of the information mentioned above are provided in the final output table of ‘local’ and ‘global’ functions. The hits cutoff we used in this study are mentioned in the main text.

#### HiC and eQTL overlapping analysis

We obtained the MCF7 (ER+) breast cancer cell line contact domain data from the ENCODE project (accession: ENCFF164AGX) as an approximation ^42^ since there is no publicly available MDA-MB-361 HiC dataset. We overlapped the 3,509 MCF7 contact domains with the 369 MDA-MB-361 local hits with ‘bedtools intersect’, and found that 103 (28%) of the perturbation regions with a local hit are within contact domains. These 103 regions represent the universe of possible overlaps. We found that 40 of these regions have primary target genes in the same contact domain.

We referenced 3 recent breast cancer eQTL studies and searched for overlapping regions in our local hits and reported eQTL regions ^43–45^. The results are reported in Supplementary Table 6.

#### Plotting

To generate the circos plots, we utilized the package pyCircos (version 0.3.0) and followed the tutorial. The detailed codes and scripts of circos plots and Manhattan plots can be found on GitHub.

## Data availability

The sequencing data generated in this study are deposited to the Gene Expression Omnibus (GEO) under the accession number: GSE223812 (ATAC-seq, reviewer token: yxyzyykytbgjbap), GSE224986 (Single-cell libraries, reviewer token: wtcbikkqpvinter) and GSE228332 (Bulk RNA-seq, reviewer token: azqdscaafpqjbgj). The epigenomic bigwig files of MDA-MB-361 and MDA-MB-231, variants bed files and sgRNA bed files were deposited under the supplementary files of GSE223812. We used these processed files to generate all the genome browser snapshots in this study. The unfiltered global hit tables for MDA-MB-361 and MDA-MB-231 have also been deposited in the GSE224986 supplementary files.

## Code availability

pySpade is available at PyPi (https://pypi.org/project/pySpade/) and on GitHub (https://github.com/Hon-lab/pySpade). Other analysis scripts and notebooks of this study are available on GitHub (https://github.com/Hon-lab/Breast_cancer_perturbation).

## AUTHOR CONTRIBUTIONS

**Yihan Wang**: Software, Validation, Formal Analysis, Investigation, Visualization, Writing-Original draft, Writing-Review & Editing. **Daniel Armendariz:** Investigation, Formal Analysis. **Lei Wang**: Investigation, Validation. **Huan Zhao**: Validation. **Shiqi Xie**: Conceptualization. **Gary Hon**: Conceptualization, Writing-Original draft, Writing-Review & Editing, Supervision, Project Administration, Funding Acquisition.

## ACKNOWLEDGEMENTS

We acknowledge the BioHPC computational infrastructure at UT Southwestern for providing HPC and storage resources that have contributed to the research results reported within this paper. We thank W. Lee Kraus and Tulip Nandu for kindly sharing the LONESTAR Consortium data. We acknowledge Hyung Bum Kim, W. Lee Kraus and Boxun Li for carefully reading the manuscript and providing suggestions to improve this study. We thank members from Hon Lab, Banaszynski Lab and Munshi Lab for helpful discussion. G.C.H is supported by CPRIT (RP190451), NIH (DP2GM128203, UM1HG011996), the Burroughs Wellcome Fund (1019804), the Welch Foundation (I-2103-20220331), and the Green Center for Reproductive Biology.

## SUPPLEMENTAL FIGURE LEGENDS

**Supplementary Figure 1:**
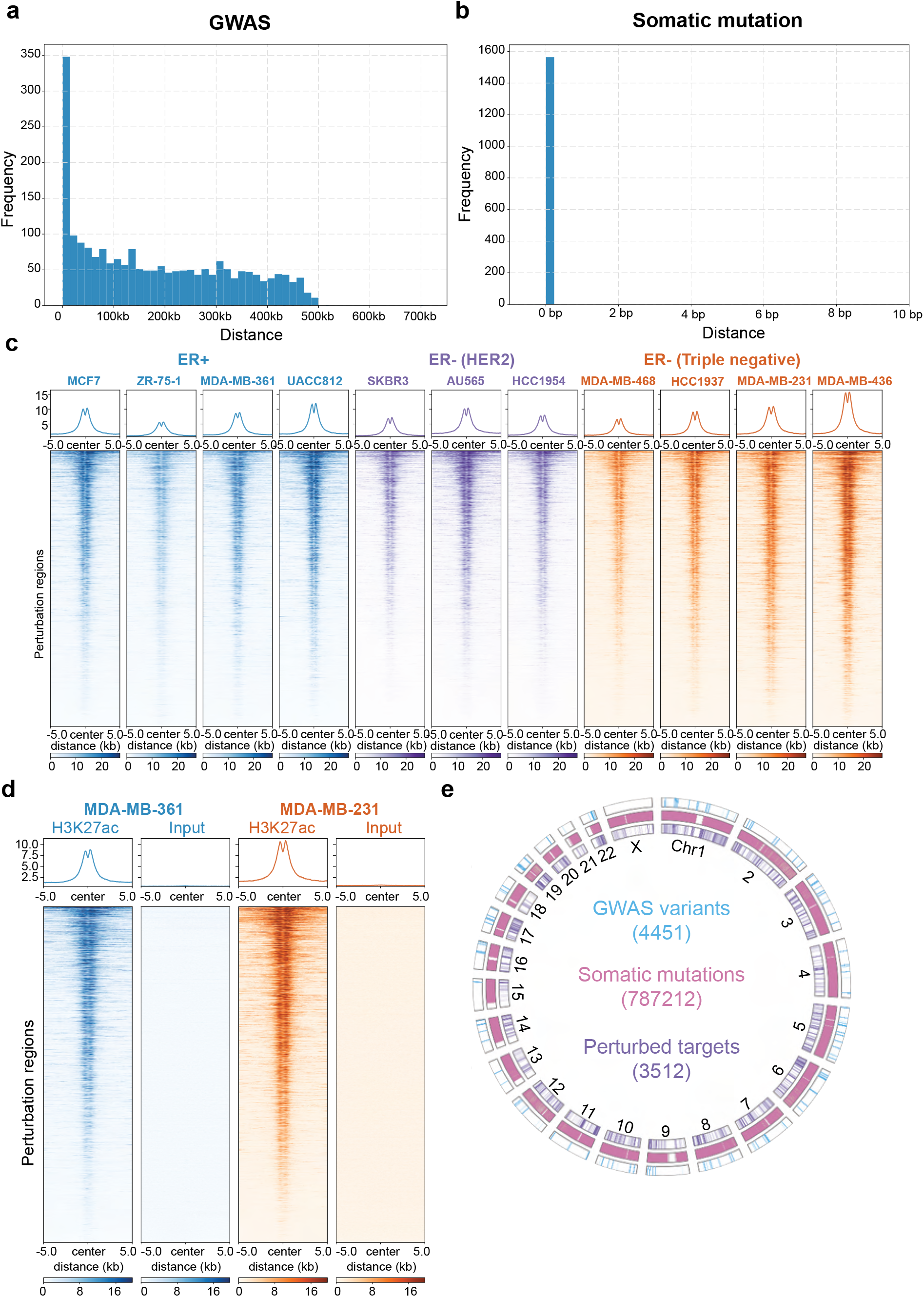
Select perturbation regions based on variants. a. The distance between GWAS SNPs and GWAS perturbation regions. b. The distance between somatic mutations and SM perturbation regions. Since the regions are selected based on overlapping with somatic mutations, the closest distances are all 0 bp. c. The H3K27ac ChIP-seq signals of 11 breast cancer cell lines in all 3,512 perturbation regions. d. The H3K27ac and input signals of (ER+) MDA-MB-361 and (ER-) MDA-MB-231 in all the perturbation regions. e. Circos plot illustrates the localization of previously identified 4,452 GWAS variants, 787,212 somatic mutations, and the 3,512 prioritized regions perturbed in this study.

**Supplementary Figure 2:**
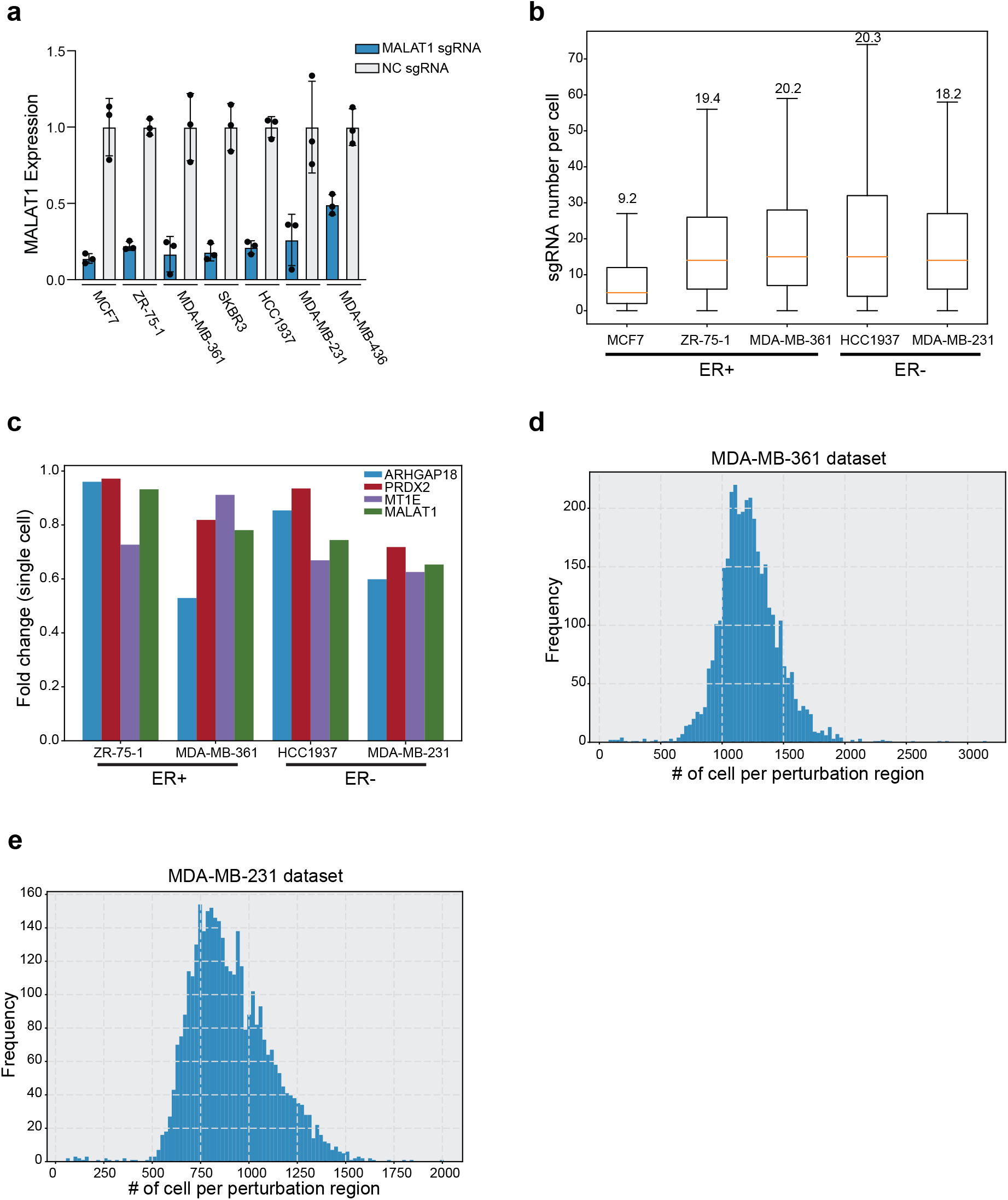
(ER+) MDA-MB-361 and (ER-) MDA-MB-231 were selected by testing different subtypes of breast cancer cell lines. a. Bulk qPCR of MALAT1 promoter in 7 breast cancer cell lines: (ER+) MCF7, (ER+) ZR-75-1, (ER+) MDA-MB-361, (ER-) SKBR3, (ER-) HCC1937, (ER-) MDA-MB-231, (ER-) MDA-MB-436 shows that dCas9-KRAB repression works in most breast cancer lines. a. b. The average number of sgRNA in each cell from single cell data. The lentiviral infectivity is low in (ER+) MCF7. b. Single-cell data of positive control promoters repression. Overall, dCas9-KRAB works better in (ER+) MDA-MB-361 and (ER-) MDA-MB-231 of each subtype. c. The distribution of the number of cells in each perturbation of MDA-MB-361 screens. The majority perturbation regions have 1,000 to 1,500 cells. d. The distribution of the number of cells in each perturbation of MDA-MB-231 screens. There are slightly less cells (750 to 1000) for each perturbation compared to the MDA-MB-361 screens.

**Supplementary Figure 3:**
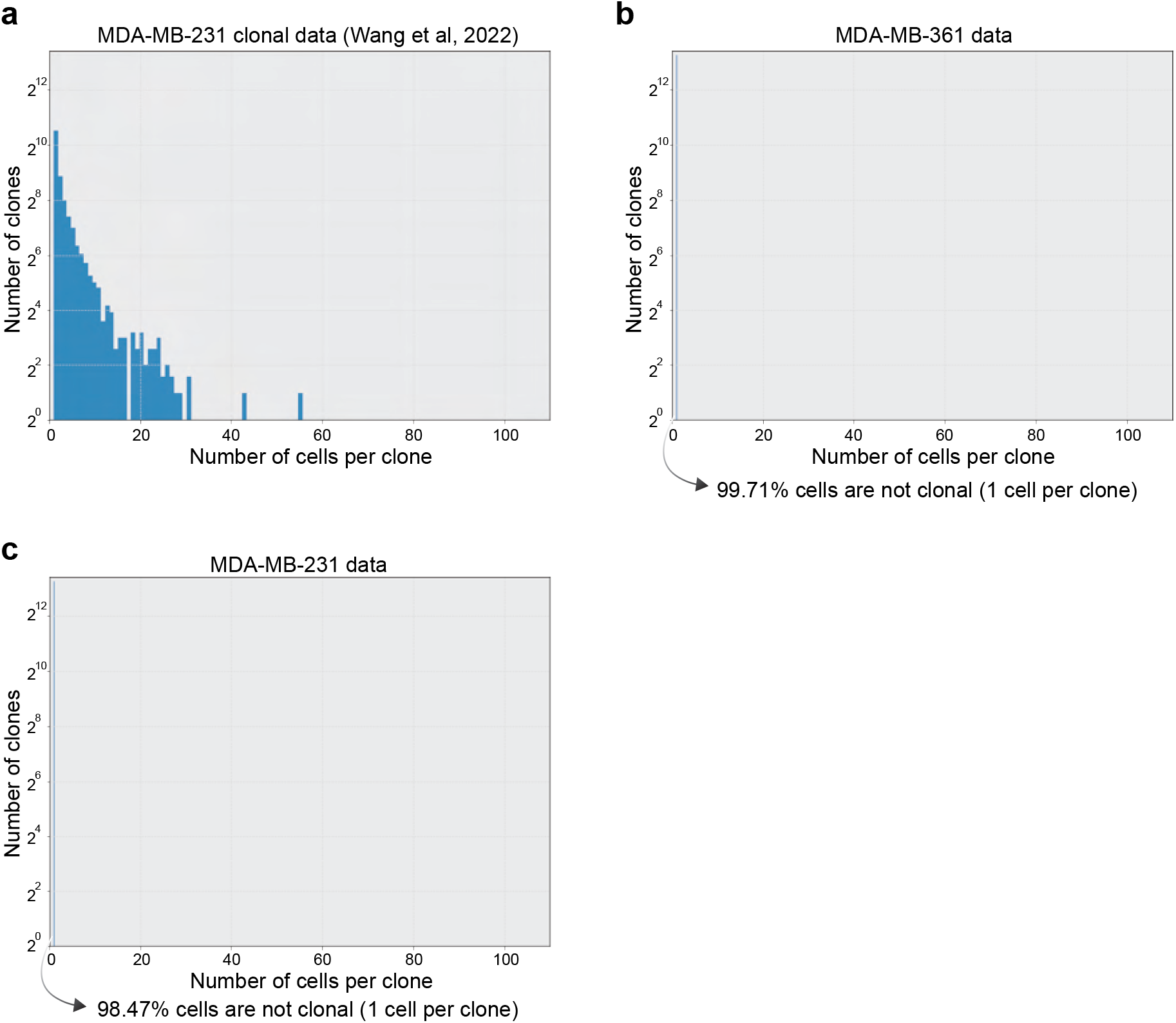
Clonality does not affect the quality of this study. a. Our previous study showed that clonality affects the ability to identify hits. Dozens of cells are found in each clone in our previous study. b. The clonal analysis of MDA-MB-361 cells of this study, indicating clonality does not affect the differential expression analysis. c. The clonal analysis of MDA-MB-231 cells of this study, indicating clonality does not affect the analysis.

**Supplementary Figure 4:**
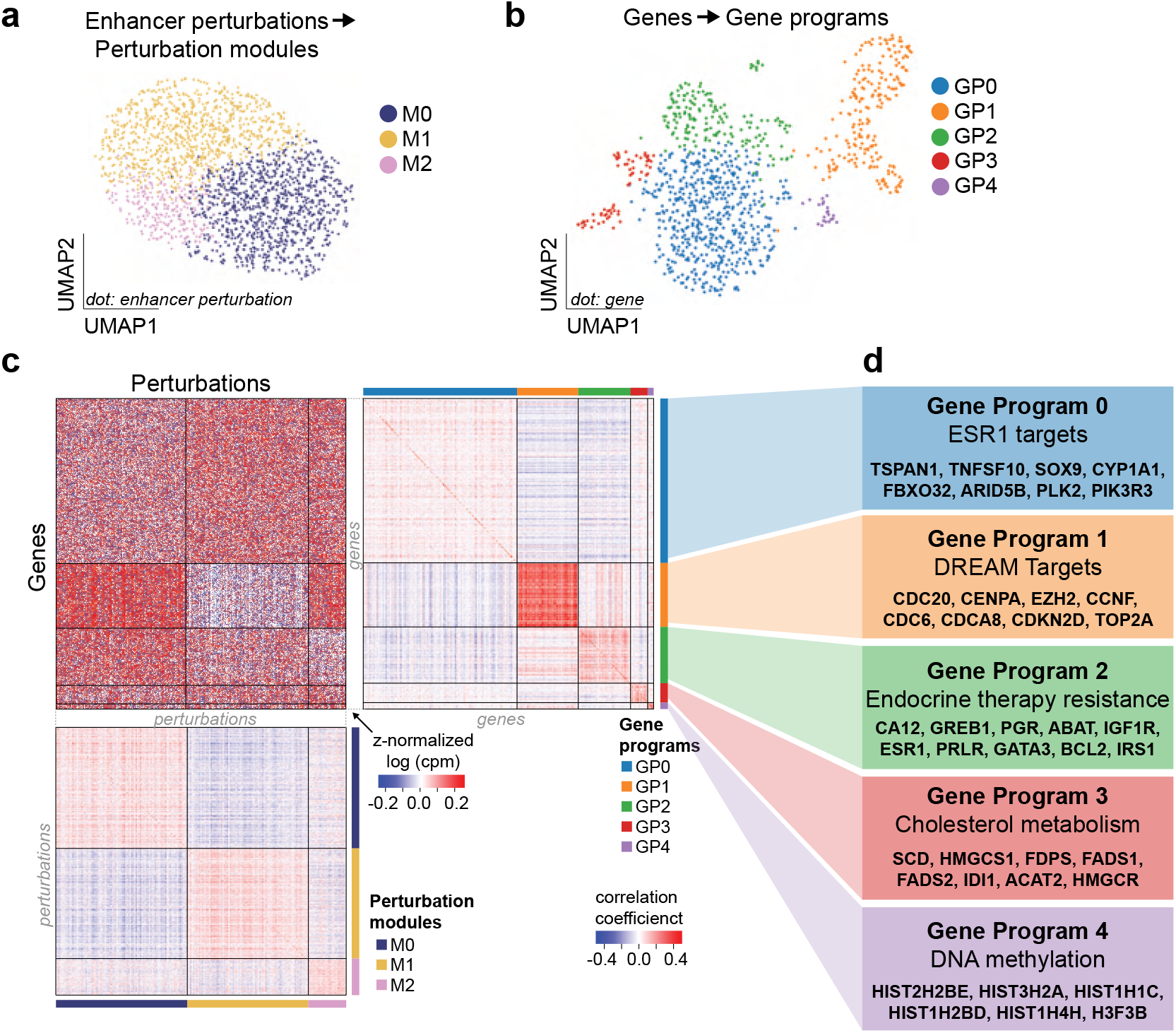
Perturbation modules and gene programs of MDA-MB-361 GWAS dataset. a. The UMAP plot of enhancer perturbations. b. The UMAP plot of gene programs. c. (Upper left) The normalized expression heatmap arranged by perturbation modules and gene programs. (Upper right) Correlation coefficient heatmap of gene programs indicates highly similarity within the same gene program. (Lower) Correlation coefficient heatmap of perturbation modules confirms that enhancers within the same module share a similar expression pattern. d. Functional annotation of each gene program identified by GSEA.

**Supplementary Figure 5:**
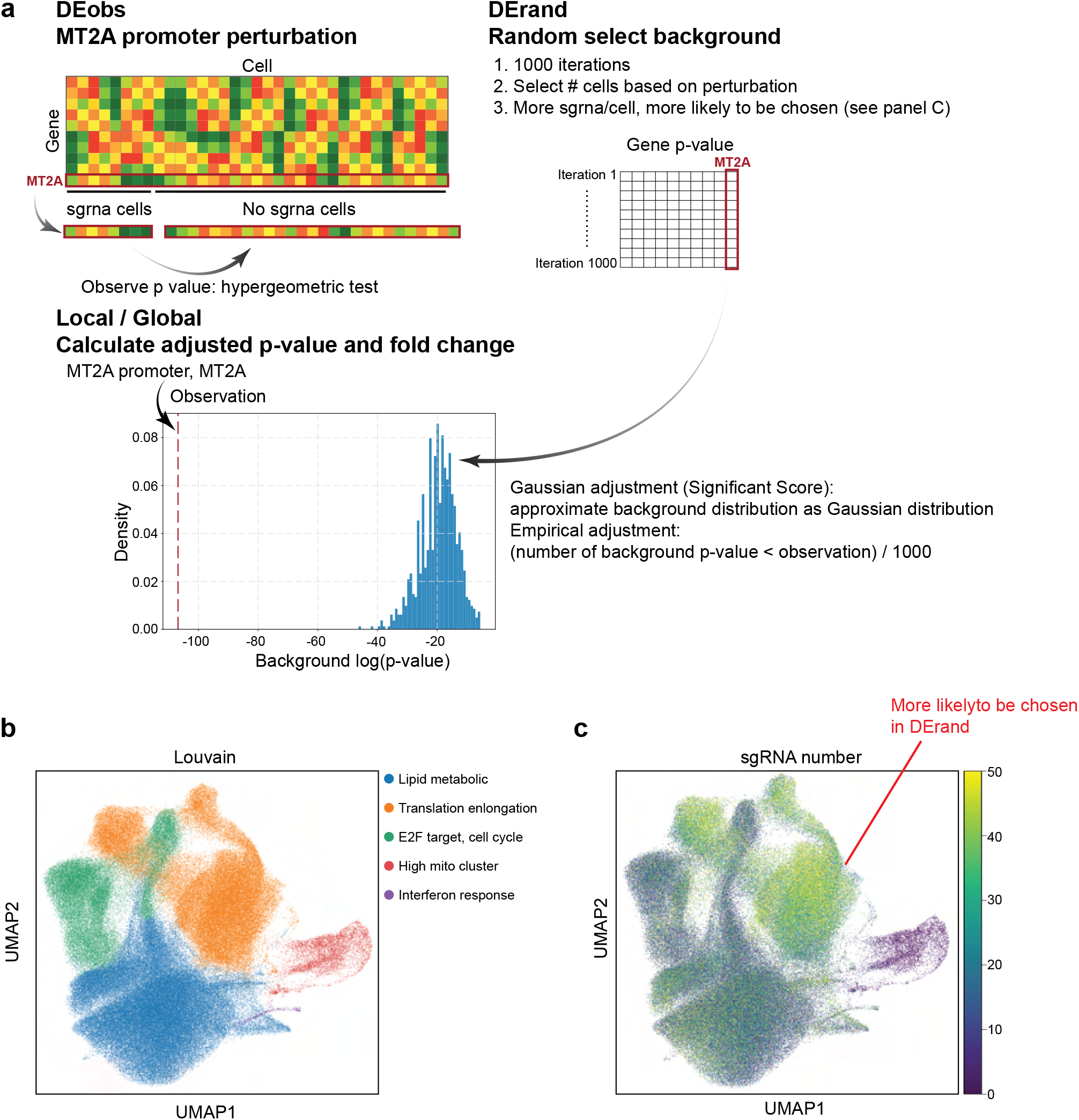
Overview of pySpade and MDA-MB-361 single-cell clustering. a. Schematic diagram of the hits calling and randomized background to calculate the empirical p-value and adjusted p-value of target genes. b. Louvain clustering of MDA-MB-361 cells shows that the transcriptome of MDA-MB-361 cell line perturbation is relatively homogenous. c. The number of sgRNA in the UMAP embedding indicates the sgRNA number is not even across different clusters.

**Supplementary Figure 6:**
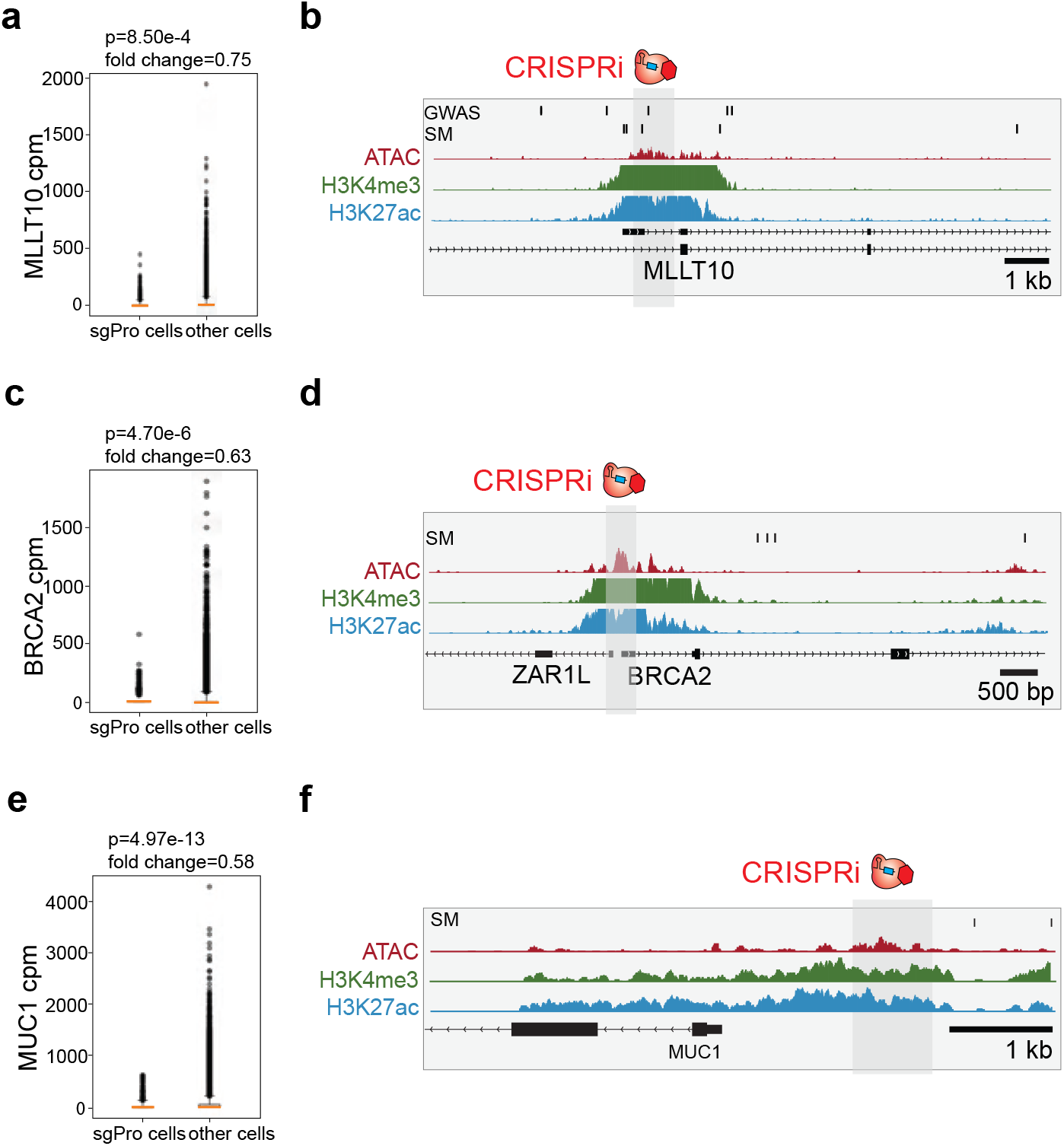
Perturb-seq identifies non-coding regulatory elements that regulate cancer genes. a. Single cell data shows perturbing MLLT10 promoter leads to MLLT10 down-regulation. (Raw p-value, Student’s t-test). (Box plot: center line, median; box limits, top and bottom 10%; whiskers, 1.5x interquartile range; points, outlier). b. Genome browser tracks of MLLT10 promoter locus. c. Single cell data shows that perturbing BRCA2 promoter causes BRCA2 loss of expression. (Raw p-value, Student’s t-test). (Box plot: center line, median; box limits, top and bottom 10%; whiskers, 1.5x interquartile range; points, outlier). d. Genome browser snapshot of BRCA2 promoter region. e. Single cell data of perturbing MUC1 promoter causing MUC1 down-regulation. (Raw p-value, Student’s t-test). (Box plot: center line, median; box limits, top and bottom 10%; whiskers, 1.5x interquartile range; points, outlier). f. Epigenetic genome tracks of MUC1 promoter locus.

**Supplementary Figure 7:**
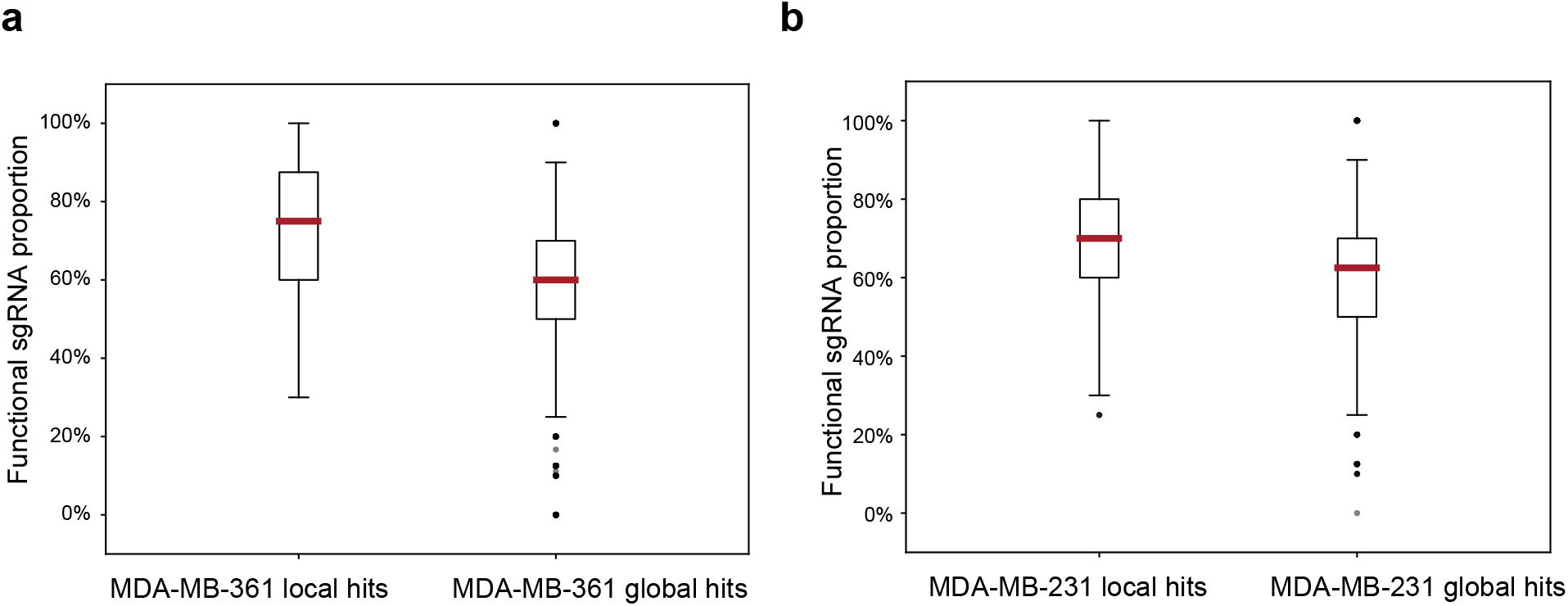
sgRNAs targeting the same region are functionally consistent with each other. a. The proportion of sgRNAs that are functional for each perturbation region. The boxes indicate the local and the global hits of MDA-MB-361 cells. For each hit, if individual sgRNA perturbation leads to the same effect as combining all the sgRNAs, and the fold change is more than 20%, we consider the sgRNA functional. (Box plot: center line, median; box limits, upper and lower quartiles; whiskers, 1.5x interquartile range; points, outlier). b. The box plots indicate that more than half of the sgRNAs for each perturbation region are functional. (Box plot: center line, median; box limits, upper and lower quartiles;; whiskers, 1.5x interquartile range; points, outlier).

**Supplementary Figure 8:**
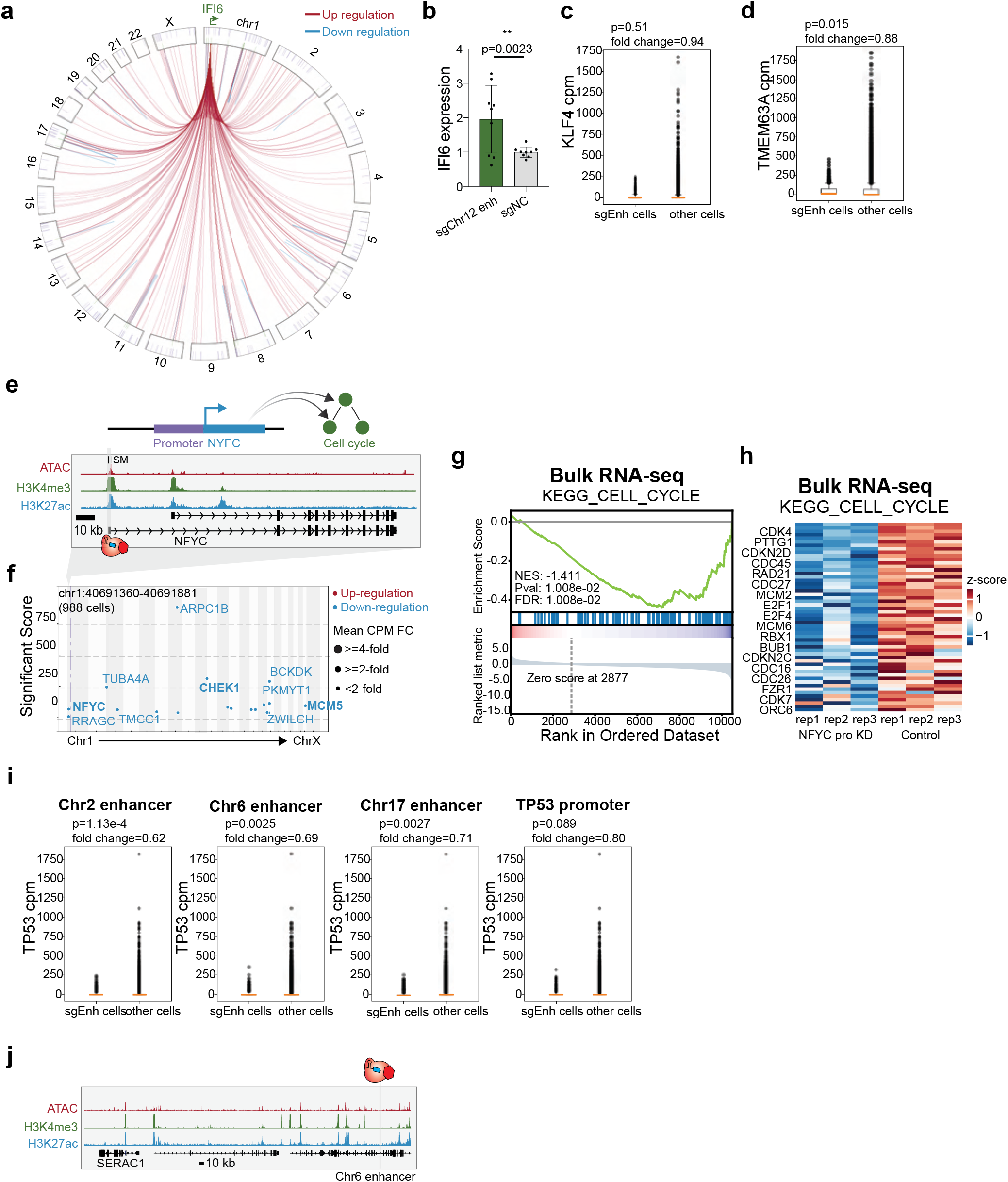
Single-cell data of enhancer indirect hits. a. The circos plot shows that multiple enhancers indirectly regulate IFI6 located on chromosome 1. b. Bulk qPCR verified one enhancer (chr12:12716861-12717861) indirectly activates IFI6. (NC: non-target control sgRNA, * p< 0.05, ** p< 0.01, *** p< 0.001, Student’s t-test). c. The single-cell data of KLF4 expression with KLF4 enhancer perturbation, which is validated by bulk qPCR experiments. (Raw p-value, Student’s t-test). (Box plot: center line, median; box limits, top and bottom 10%; whiskers, 1.5x interquartile range; points, outlier). d. The single-cell data of TMEM63A expression with chr1 enhancer knock-down. (Raw p-value, Student’s t-test). (Box plot: center line, median; box limits, top and bottom 10%; whiskers, 1.5x interquartile range; points, outlier). e. Genome browser snapshot of the NYFC locus, with targeted region indicated. f. Manhattan plot of NYFC promoter perturbation shows that multiple cell cycle genes are down-regulated. The dotted line indicates the perturbed region. g. Gene Set Enrichment Analysis (GSEA) of bulk RNA-seq validation experiment shows that the cell cycle pathway is down-regulated with NFYC perturbation. h. The heatmap derived from bulk RNA-seq analysis shows that multiple genes of the cell cycle pathway are down-regulated with NFYC promoter knock-down. i. TP53 single-cell expression data with three indirect enhancers (chr2:201258117-201258617, chr6:158737687-158738187 and chr17:39178360-39178946) and the TP53 promoter (chr17:7687427-7688427) knock-down. (Raw p-value, Student’s t-test). (Box plot: center line, median; box limits, top and bottom 10%; whiskers, 1.5x interquartile range; points, outlier). j. Genome browser snapshot of chr6 enhancer that indirectly regulates TP53 through SERAC1.

**Supplementary Figure 9:**
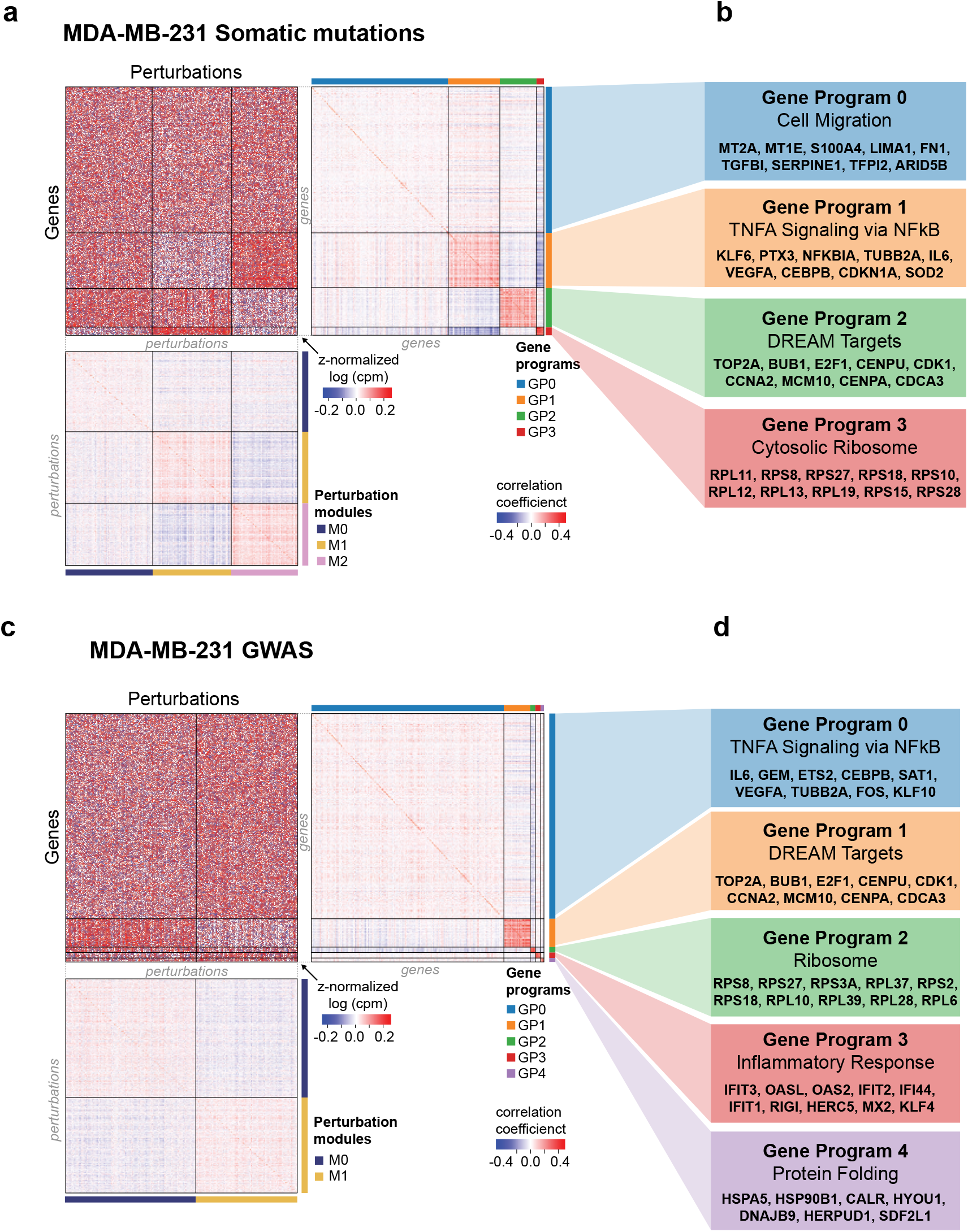
Perturbation modules and gene programs of MDA-MB-231 screens. a. The heatmaps of perturbation modules and gene programs of MDA-MB-231 cells somatic mutations screen. b. Functional annotations of each gene program. c. The heatmaps of perturbation modules and gene programs of MDA-MB-231 cells GWAS screen. d. Functional annotations of each gene program.

**Supplementary Figure 10:**
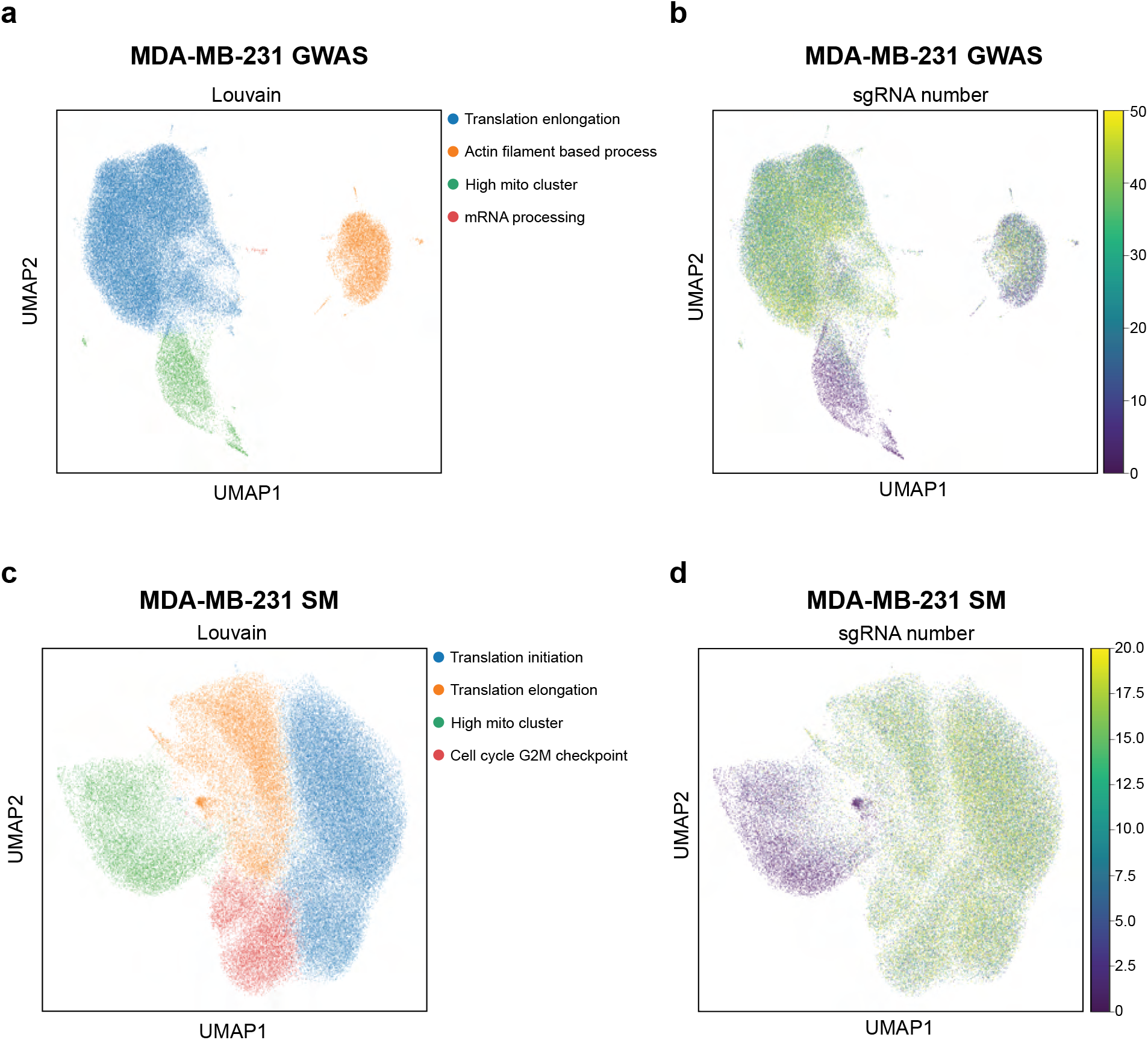
MDA-MB-231 single-cell clustering. a. Louvain clustering of MDA-MB-231 cells with GWAS perturbation screen. b. The number of sgRNA in UMAP embedding in the MDA-MB-231 GWAS dataset. c. Louvain clustering of MDA-MB-231 cells with somatic mutations (SM) screen. d. The number of sgRNA in UMAP embedding in the MDA-MB-231 SM dataset.

**Supplementary Figure 11:**
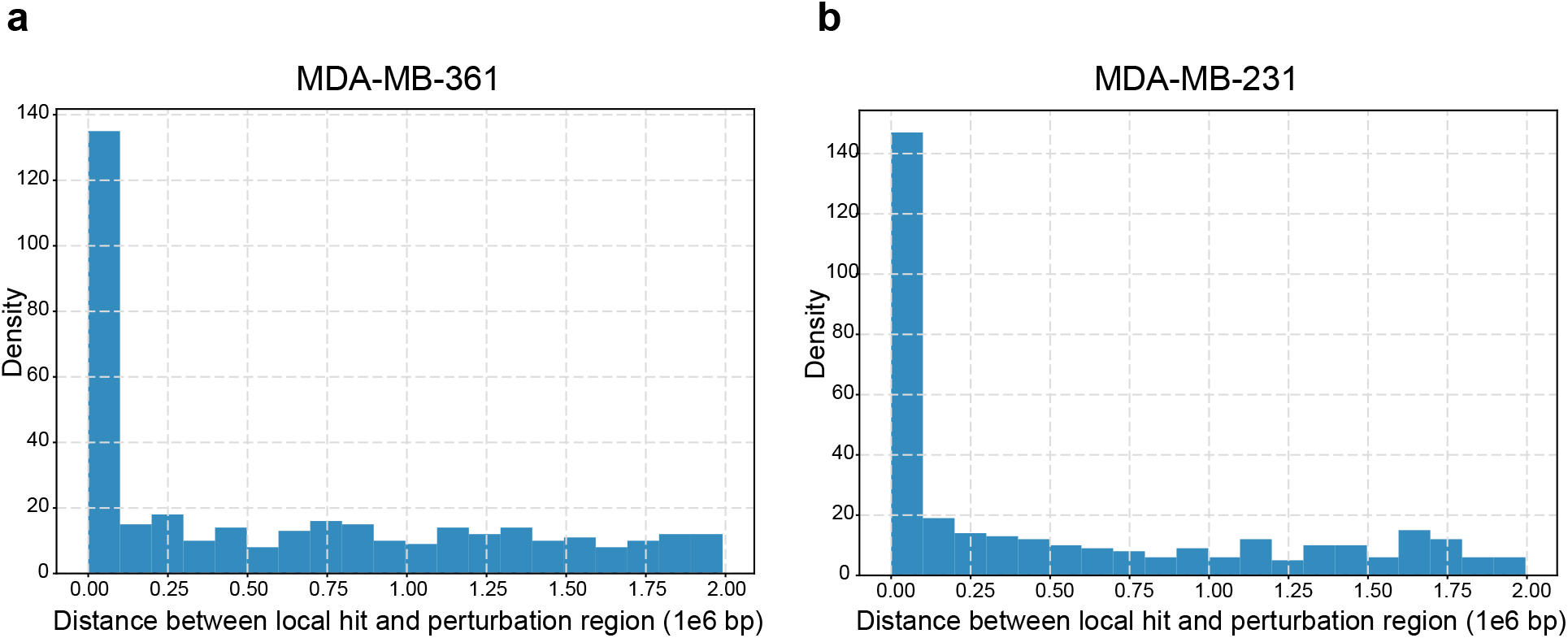
Local hit comparison in ER+ and ER- cells. a. The distance between the perturbation region and local hit genes in (ER+) MDA-MB-361 cells. b. The distance between the perturbation regions and local hit genes in (ER-) MDA-MB-231 cells.

**Supplementary Figure 12:**
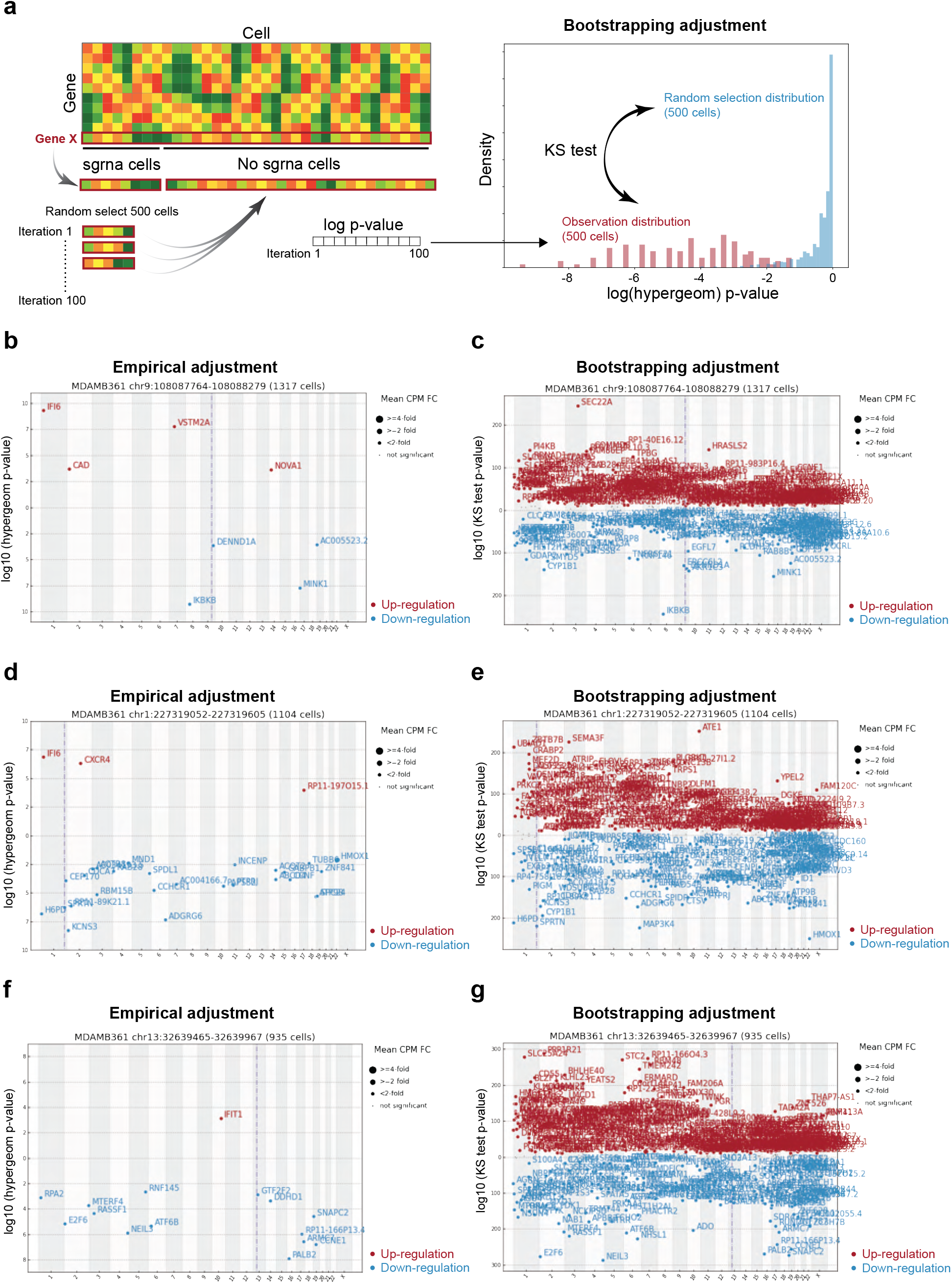
Comparison between empirical adjustment and bootstrapping adjustment. a. Schematic illustration of bootstrapping adjustment method. b. Manhattan plot of IKBKB enhancer with empirical p-value cutoff hits. The y axis indicates the raw hypergeometric test p-value. The Gaussian adjustment Manhattan plot for IKBKB enhancer is shown in Fig. 3d. c. Manhattan plot of IKBKB enhancer with bootstrapping adjustment. d. Manhattan plot of HMOX1 enhancer with empirical p-value cutoff hits. The Gaussian adjustment Manhattan plot for HMOX1 enhancer is shown in Fig. 3j. e. Manhattan plot of HMOX1 enhancer with bootstrapping adjustment. f. Manhattan plot of chr13 enhancer with empirical p-value cutoff hits. The Gaussian adjustment Manhattan plot for chr13 enhancer is shown in Fig. 4d. g. Manhattan plot of chr13 enhancer with bootstrapping adjustment.

